# Engineered molecular sensors of cell surface crowding

**DOI:** 10.1101/2022.11.18.517164

**Authors:** Sho C. Takatori, Sungmin Son, Daniel Lee, Daniel A. Fletcher

## Abstract

Cells mediate interactions with the extracellular environment through a crowded assembly of transmembrane proteins, glycoproteins and glycolipids on their plasma membrane. The extent to which surface crowding modulates the biophysical interactions of ligands, receptors, and other macromolecules is poorly understood due to the lack of methods to quantify surface crowding on native cell membranes. In this work, we demonstrate that physical crowding on reconstituted membranes and live cell surfaces attenuates the effective binding affinity of macromolecules such as IgG antibodies in a surface crowding-dependent manner. We combine experiment and simulation to design a crowding sensor based on this principle that provides a quantitative readout of cell surface crowding. Our measurements reveal that surface crowding decreases IgG antibody binding by 2-20 fold in live cells compared to a bare membrane surface, resulting in a cell surface osmotic pressure opposing binding of 1 - 4 kPa. Our sensors show that sialic acid, a negatively charged monosaccharide, contributes disproportionately to red blood cell surface crowding via electrostatic repulsion, despite occupying only ~1% of the total cell membrane by mass. We also observe significant differences in surface crowding for different cell types and find that expression of single oncogenes can both increase and decrease crowding, suggesting that surface crowding may be an indicator of both cell type and state. Our high-throughput, single-cell measurement of cell surface osmotic pressure may be combined with functional assays to enable further biophysical dissection of the cell surfaceome.

**Significance Statement:** Cells interact with each other and the extracellular environment through a crowded assembly of polymers on their plasma membranes. The high density of these surface polymers can generate physical crowding that impacts cell function. However, tools to quantify the extent and effect of surface crowding on live cell membranes are lacking. In this work, we design macromolecular sensors that act as direct reporters of cell surface crowding. We combine experiments on reconstituted and live cell surfaces with molecular dynamics simulations to provide a mechanistic understanding of how cell surface crowding reduces binding of soluble molecules, and we show that crowding varies significantly with cell type and is affected by oncogene expression.

## Introduction

The biophysical organization of proteins, glycoproteins, and glycolipids that densely coat the surface of a cell’s plasma membrane has been shown to govern many important physiological processes. Physical crowding on cell surfaces has a direct connection to cancer malignancy (1–3), and the dense glycocalyx on cancer cell surfaces can sterically hinder antibody binding and phagocytosis by immune cells (4). Recent studies have shown that the glycocalyx can also attenuate the binding of viruses and lectins to cell surface receptors (5–7). Surface crowding also alters protein mobility and sorting (8), as well as membrane channel gating (9). However, quantitative methods to obtain a detailed, mechanistic understanding of plasma membrane density and the biophysical interactions that govern macromolecular binding on live cell surfaces are lacking. This leaves basic questions about cell surface crowding unanswered, including the extent to which glycosylation contributes, how crowding differs among cell types and states, and even how best to quantify crowding in live cells.

While proteomic analysis of cell surface proteins provide detailed information on the relative abundance of proteins at the population level (10), it is difficult to predict collective biophysical features of cell surfaces simply from knowledge of the surface proteome. Previous studies have shown that the physical accessibility of large soluble ligands and macromolecules decreases on synthetic surfaces grafted with synthetic polymers or purified proteins (11–14). Other studies have developed tools that measure effects of surface crowding, including Houser et al. (15), who measured the separation of FRET pairs as a function of the steric interactions within the surface polymers on reconstituted membranes, and Son et al. (16), who measured nanometer-scale changes in height of multi-domain proteins *in vitro* as surface density increased. While these studies of reconstituted systems provide valuable insights, direct quantification of surface crowding on live cell membranes remains a challenge. Advanced imaging techniques like electron microscopy enable nanometer-scale visualization of the cell membrane (17), but structural information does not easily translate to a physical understanding of the effects of crowding, leaving a need for new tools to study the surfaces of living cells (3).

Here we report a simple and direct approach to measure surface crowding on live cells. Inspired by theoretical work on the adsorption of macromolecules on crowded surfaces (18), we engineered macromolecular probes that insert into bilayer membranes and directly quantify the repulsive penalty posed by crowded cell surfaces by a reduction in effective affinity. We first validated the measurement principle by adding polymer-cholesterol conjugates of varying size to reconstituted membranes or red blood cells and analyzing the result with molecular dynamics (MD) simulations and adsorption theories. We then designed a two-component crowding sensor based on a biotin-fluorophore-cholesterol conjugate that spontaneously incorporates into live cell plasma membranes and a monomeric anti-biotin antibody introduced in solution around the cell. By measuring the fraction of antibody bound to the surface biotin in individual cells relative to bare model membranes, we can directly quantify the effect of cell surface crowding on antibody binding affinity. Our crowding sensors work with cells of different size, shape, and membrane lipid compositions, and they enable absolute quantification of surface crowding in terms of an osmotic pressure, providing a way of comparing across different cell types and states.

## Results

### Physical crowding on membrane surfaces reduces binding affinity of soluble macromolecules

Existing theories of adsorption thermodynamics give a direct relationship between the dissociation constant of a soluble macromolecule and the free energy of the surface, *K_D_* = exp(*U*/(*k_B_T*)) (Fig. 1A). Therefore, a measurement of the effective *K_D_* on a crowded surface is a direct reporter of the free energy barrier posed by the crowded surface. To explore whether we could use this principle to directly read out surface crowding, we conducted coarse-grained molecular dynamics (MD) simulations to study the binding of macromolecules to an exposed surface and to a surface decorated with polymers (Fig. 1A). Based on simulations, the free energy as a function of distance from the surface increases in the presence of surface-tethered polymers (Fig. 1B), and the deviation of the energy minima is a direct readout of the effective dissociation constant, *K_D_* (Supplemental Fig. 1; see also Materials and Methods). As expected, we find that *K_D_* increases as we increase the surface crowding by changing either the surface density or the contour length of the crowding polymers (Fig. 1C). It is important to note that the contour length of the surface polymers has a strong effect on *K_D_*, which demonstrates that the polymer number density alone is an insufficient metric of surface crowding.

**Figure 1.**
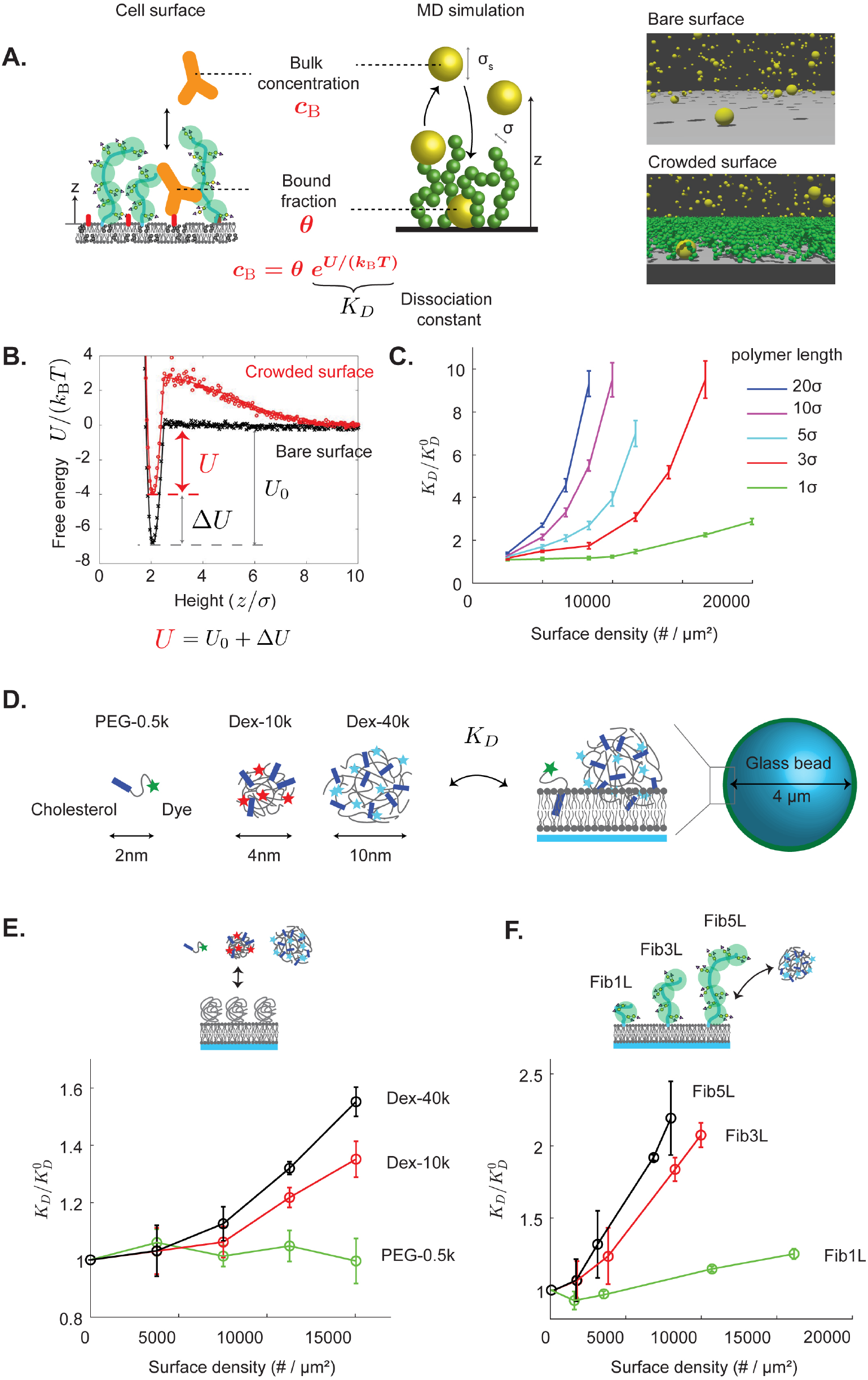
Macromolecular binding is a direct reporter of surface crowding. (A) Soluble macromolecules, like monoclonal antibodies, experience a repulsive energy penalty when binding to antigen targets located on crowded surfaces. Therefore, the concentration of surface-bound macromolecules is a direct readout of the effective dissociation constant and the crowding state of the surface. (B) Coarse-grained molecular dynamics (MD) simulations are used to calculate the binding energy of soluble macromolecules on bare (black symbols) and crowded (red symbols) surfaces. The binding affinity of soluble macromolecules on crowded surfaces (*U*) is smaller than that on bare surfaces (*U*_0_). The difference, Δ*U* = *U* – *U*_0_, is directly related to the crowding-induced change in the effective dissociation constant, *K_D_*. Polymer brush theories are used to relate the effective dissociation constant to surface crowding (solid curves). (C) Snapshots of MD simulations on bare (top) and crowded (bottom) surfaces. Soluble macromolecules (yellow spheres) of size *σ_s_* bind onto a surface functionalized with multi-domain protein polymers (green spheres) with density, *n*, and monomer size, *σ*. The effective dissociation constant, *K_D_*, normalized by the bare-surface value, 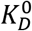, increases monotonically with the surface protein density and contour length, *L*. See Supplementary Fig. 1 and Supplementary Information for further theoretical and computational results. (D) Experimental design of cholesterol-based sensors to measure crowding on membranes. Synthetic polymers of different molecular weights are used to vary the overall size of the sensor. The sensors have a strong binding affinity to the lipid bilayer, and fluorescent labels provide a direct optical readout of bound surface concentration and the effective dissociation constant, *K_D_*. (E) Normalized dissociation constant, 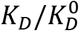, of the small (PEG-0.5k), medium (dex-10k), and large (dex-40k) sensors increases on lipid-coated beads functionalized with PEG-3k at varying surface densities. (F) 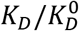 of large sensors increases on lipid-coated beads functionalized with engineered proteins of the FNIII domain repeats, Fibcon (Fib), which has a size of ~4 nm per domain. Fib1L contains 1 domain; Fib3L contains 3 domains; Fib5L contains 5 domains. For all data, error bars indicate standard deviations of the mean; N > 3.

Guided by these predictions, we engineered a series of exogenous macromolecular probes with known size and affinity to a lipid bilayer by conjugating cholesterol and fluorescent dyes to PEG 1k, dextran 10k, and dextran 40k macromolecules (Fig. 1D, see also Materials and Methods). The diameters of these probes are approximately 2, 4, and 10 nm based on the radii of gyration (19). We hypothesized that reduced incorporation of these “crowding sensors” to crowded membrane surfaces would act as a direct reporter of the energy penalty posed by surface crowding. We first tested the binding of our sensors to a supported lipid bilayer (SLB) formed on the surface of a glass bead and decorated it with PEG3k polymers simulating cell surface crowding. After allowing the sensor binding to reach equilibrium (45 minutes), we quantified the bead fluorescence with a flow cytometer. Using an adsorption isotherm to relate the bound sensor concentration to the bulk concentration, we calculated the effective dissociation constant. We found that the *K_D_* of the larger dextran 10k and 40k sensors increased on the PEG3k surface compared to that on the bare surface, with the *K_D_* of the dextran 40k sensor increasing by 55% when the SLB contained 1% (mol/mol) PEG3k (~15,000/μm^2^ area density) (Fig. 1E), which agrees with our simulation and is consistent with previous studies (12). In contrast, the small PEG 0.5k sensor experienced no change in *K_D_* for the PEG3k surface densities we tested (Fig. 1E). We observed no systematic change in the time to reach equilibrium on PEG3k surfaces compared to bare surfaces (Supplemental Fig. 2), confirming that none of our sensors are transport-limited on the surface.

To test the sensors’ ability to read out crowding due to proteins rather than synthetic PEG polymers, we then reconstituted SLBs with engineered proteins based on repeats of the FNIII domain, Fibcon, which has a size of ~4 nm per domain (20). We used different lengths based on one (Fib1L), three (Fib3L), or five (Fib5L) domain proteins with deca-histidine tags that bind to SLBs containing 1,2-dioleoyl-sn-glycero-3-(N-(5-amino-1-carboxypentyl)iminodiacetic acid)succinyl (DGS-NTA). The *K_D_* of the dextran 40k sensor increased by more than 100% on surfaces crowded with Fib3L and Fib5L at a density of ~10,000/μm^2^ relative to a bare membrane, whereas the *K_D_* increased by only ~20% for Fib1L (Fig. 1F). It is important to note that the molecular weight of Fib1L (~14 kDa) is 4.6 times larger than PEG3k, yet the crowding strength generated by the PEG surface is 25% larger when comparing them at the same number density. These results highlight the fact that the molecular weight of a surface species is not a proper metric of crowding, just as number density and height of surface species are insufficient metrics by themselves. Indeed, ‘crowding’ as it affects the affinity of soluble molecules at the cell surface is a collective phenomenon that includes these and other cell surface molecular properties.

The results of our simulation and reconstituted SLB experiments demonstrate the direct link between surface crowding and effective binding affinities of large macromolecules. Furthermore, we show with experiments and simulations that a simple measurement of *K_D_* is a quantitative reporter of surface crowding, regardless of the chemical identity of the surface species. The sensitivity of our sensors to protein length, molecular weight, and density provides a unique approach for studying the biophysical features of the cell surface.

### Macromolecular binding at cell surfaces is osmotically regulated by sialylation

We next used the crowding sensors to investigate the effect of glycosylation on crowding, both *in vitro* and on native cell membranes. We reconstituted lipid-coated beads with the extracellular domain of Glycophorin A (GYPA), a single-pass, mucin-like transmembrane protein with a heavily glycosylated and sialylated chain (21). We found that the effective *K_D_* of our dextran 40k sensor increased by 2x on the crowded GYPA surface compared to a bare membrane (Fig. 2A). When we treated the GYPA-coated beads with sialidase from Clostridium perfringens (C. welchii), we found that sialic acid removal decreased *K_D_* by 60% compared to the untreated GYPA surfaces. We hypothesized that the negative charge on sialic acids swells the GYPA chains, analogous to a swollen polyelectrolyte polymer brush (22), reducing the accessibility of the sensors to the membrane surface. It is important to note that the number density of GYPA on the bead surface is identical with and without sialidase treatment.

**Figure 2.**
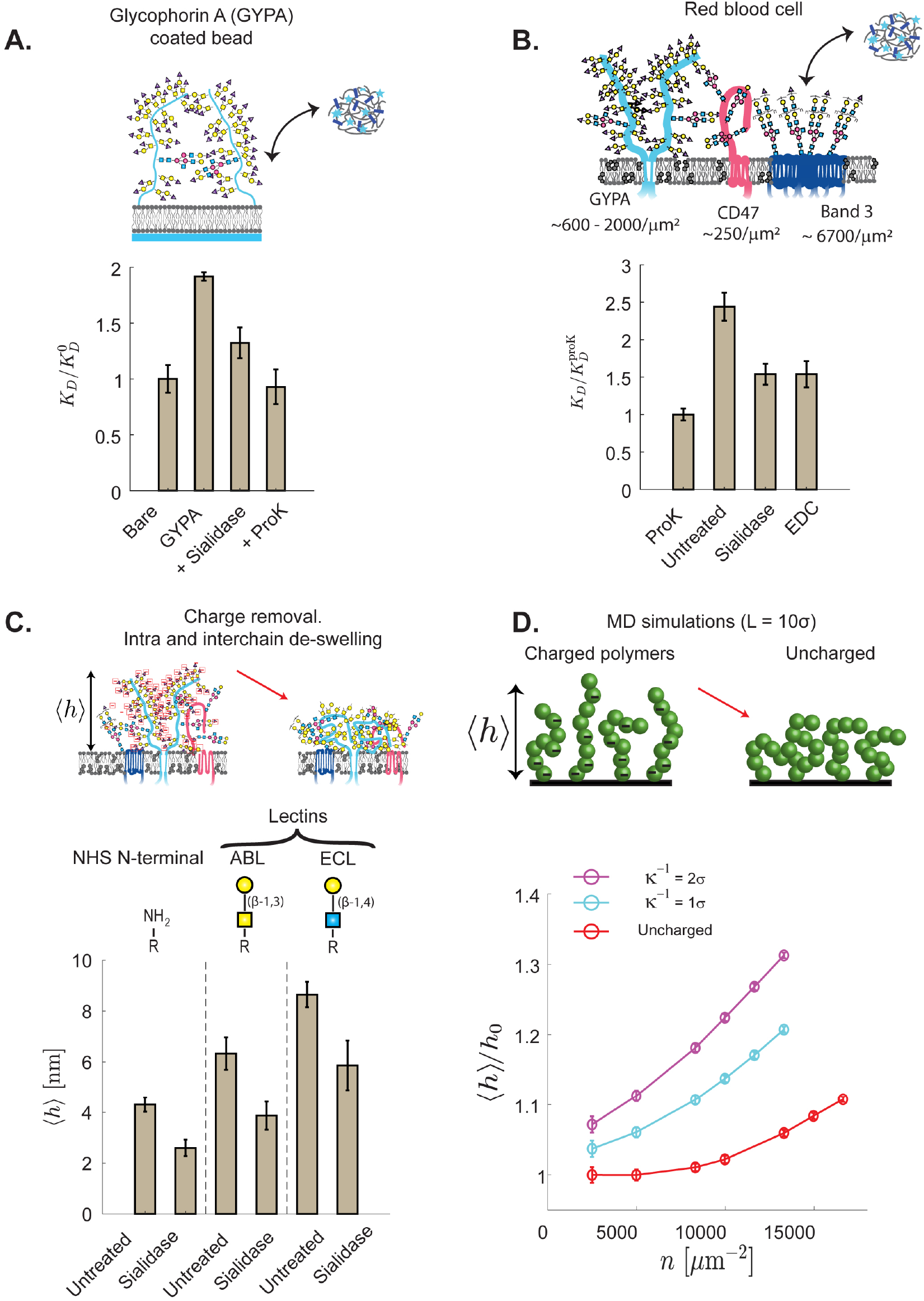
Macromolecular binding on red blood cell surfaces is osmotically regulated by sialylation. (A) Normalized dissociation constant, 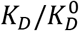, on reconstituted membranes with Glycophorin A (GYPA), a mucin-like glycoprotein found on red blood cells (RBCs). Treatment of the GYPA-bound beads with sialidase removes the sialic acids, which de-swells the GYPA and increases the surface accessibility to the sensor. Proteinase K (proK) removes GYPA and the affinity is restored to the same value as bare beads. (B) Dissociation constant on live RBCs, normalized by the affinity on proK-treated cells, 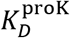. Sialidase treatment removes the sialic acids on the RBC surface and decreases *K_D_*, consistent with the results on GYPA-bound beads. EDC carbodiimide chemistry removes the negative charge on the RBC surface. The *K_D_* reduction is the same as that with sialidase treatment, consistent with our hypothesis that electrostatic charge repulsion within the glycocalyx reduces the accessibility of the sensors. (C) Average height of the RBC surface proteins, 〈*h*〈, is measured in an untreated cell and the cells treated with sialidase using the cell surface optical profilometer (CSOP) (16). The reduction in 〈*h*〉 with sialidase treatment is consistent with the de-swelling of polyelectrolyte brushes from charge removal. Measurements of RBC surface heights are based on N-terminal labeling of proteins, FITC-conjugated ABL targeting the Gal-GalNac disaccharide, FITC-conjugated ECL targeting the Gal-GlcNac disaccharide. (D) Average normalized height of multi-domain proteins in MD simulations with uncharged (red circles) and charged (cyan and magenta circles) protein residues, where *h*_0_ is the uncharged protein height at dilute densities. Electrostatic interactions among the proteins are modeled by a Yukawa potential with different Debye lengths, *κ*^-1^. The simulated decrease in surface protein height upon charge removal is consistent with the CSOP data. For all data, error bars indicate standard deviations of the mean; N > 3.

To study the extent of crowding posed by sialylation on live cell membranes, we examined human red blood cell (RBC) membranes, where the average protein compositions and copy numbers are well characterized (21,23). Approximately 23% of the RBC membrane surface area is occupied by proteins (24), with GYPA and Band 3 being two of the bulkiest and most abundant proteins. Anionic transporter Band 3 is a multi-pass transmembrane protein with a single extracellular N-glycan with poly-LacNac glycans. Given a RBC surface area of 150 *μm*^2^, we estimate a surface density of 6700/*μm*^2^ for Band 3 (5 × 10^5^ - 10^6^ copies per cell) and 1300/*μm*^2^ for GYPA (1 × 10^5^ - 3 × 10^5^ per cell) (23,25).

We found that the *K_D_* of the dextran 40k sensor is ~2.5 times larger on the RBC surface compared to that on RBC surfaces treated with broad-spectrum serine protease, proteinase K (ProK) (Fig. 3A). We note that ProK treatment leads to only a partial digestion of the cell surface proteins, so the relative crowding state on the untreated cells are even larger when compared against bare lipid-coated beads (discussed below). To study the role of surface charges, we treated RBCs with sialidase and found that the *K_D_* decreased by ~50% compared to the untreated RBC. This might be considered surprising given that sialic acids account for only 2.4 % of the total mass of glycoproteins on a RBC surface (26). To verify that charge removal is the dominant mechanism of decreased *K_D_*, we neutralized the negative charges on the carboxylic group of sialic acids using EDC chemistry (Materials and Methods). Consistent with our hypothesis, we found that charge removal reduced *K_D_* to a value comparable to the cells treated with sialidase (Fig. 2B). This further confirms that surface crowding cannot be described by surface density nor the protein molecular mass alone, and it shows that molecular charge can be a major contributor to crowding.

**Figure 3.**
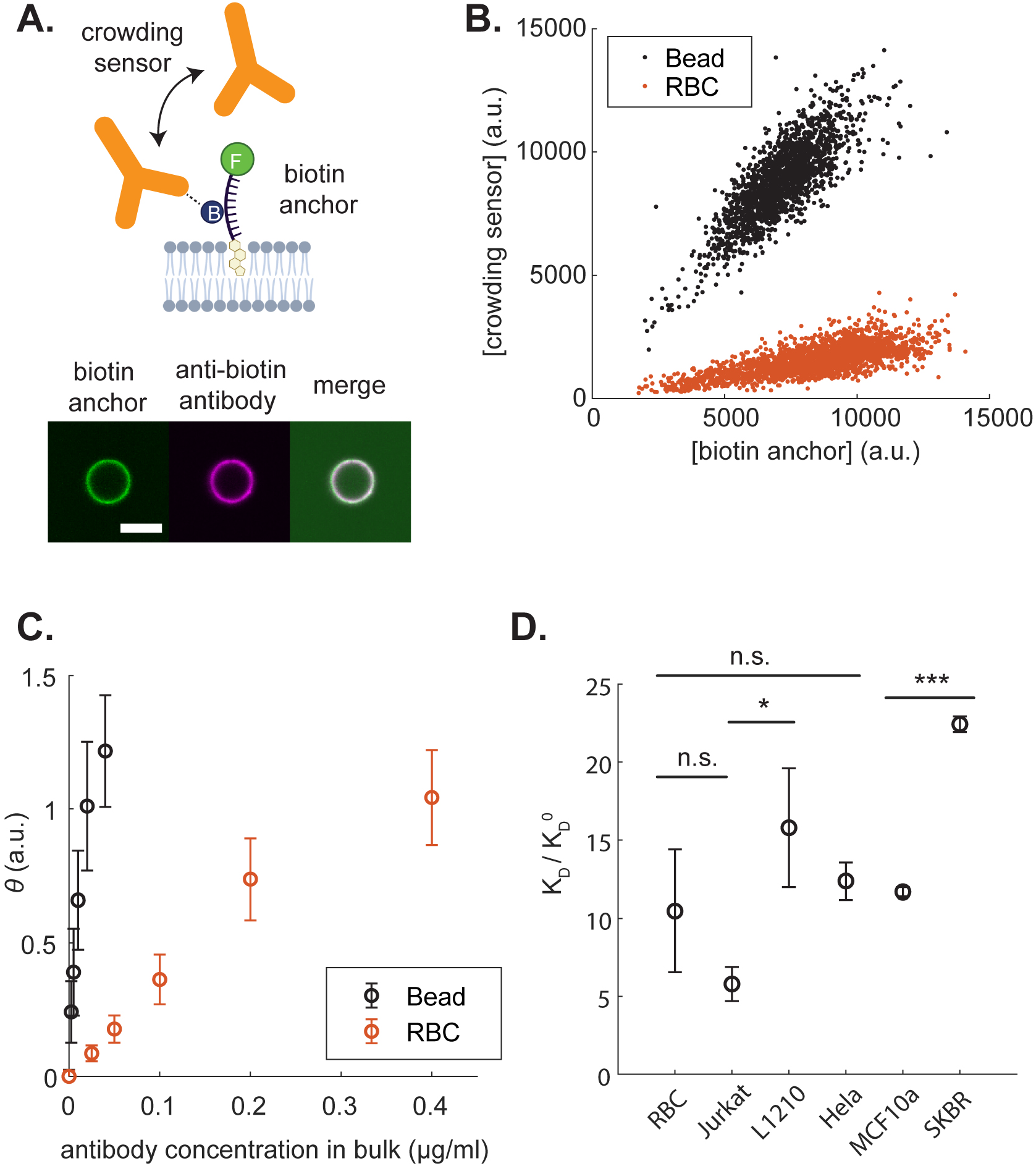
Cell surface crowding is significant and varies across different cell types. (A) Above: Schematic of the two-part crowding sensors that insert into the lipid bilayer with a biotin antigen presented above the membrane. The effective affinity of anti-biotin antibody to the biotin anchor is used as a direct reporter of surface crowding. Below: Florescence image of a lipid-coated bead with the FITC-biotin-cholesterol construct, antibiotin antibody, and the merge. Scale bare is 5 μm. (B) The amount of surface biotin anchor and the crowding sensor bound on individual RBCs or beads, measured with flow cytometry. (C) Surface concentration of bound antibody as a function of bulk concentration on a bare lipid-coated bead and RBC cell surface. The slope at small antibody concentrations is used to determine the effective binding affinity of the antibody on each surface. Error bars represent the standard deviation of *θ* within the same sample. (D) Normalized dissociation constant of antibody, 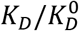, on live suspension and adherent cells. Results are normalized by the binding affinity on bare lipid-coated beads, 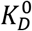. Error bars represent the standard deviation of the mean in three replicate measurements, except for Hela and SKBR, which were measured twice. P-values are calculated based on a two-tailed *Student’s* t-test (n.s.: nonsignificant, * < 0.05, ** < 0.01, *** < 0.001).

To understand the interplay between charge and glycocalyx architecture, we compared the average thickness of the RBC glycocalyx on untreated and sialidase-treated RBCs. We used Cell Surface Optical Profilometry (CSOP) (16) to measure glycocalyx thickness by quantifying the height of a fluorophore conjugated to the N-terminus of surface proteins and fluorescent lectins attached to surface glycans (Fig. 2C). We found that the average heights of both proteins and glycans reduced by ~30-40% with sialidase treatment, consistent with the notion that the polyelectrolyte brushes de-swell upon charge removal (22,27). We tested the effects of charges *in-silico* by using MD simulations of surface-tethered polyelectrolytes interacting via a screened Coulomb (Yukawa) potential with two different electrostatic interaction distances, 1.0 and 2.0 nm. We found that a surface containing mucin-like glycoproteins swells by ~40% at surface polymer densities of ~15,000 /*μm^2^* (Fig 2D), which is consistent with surface protein densities on cell membranes (21,28). Our simulations support the hypothesis that the glycocalyx maintains a swollen architecture from intra-chain and inter-chain electrostatic repulsion, and that the charged glycans pose a significant energy barrier against ligand binding. The mammalian cell surface contains approximately ~10^5^-10^6^ sialic acids / μm^2^, (1,29,30) which is further elevated in cancers (31,32), supporting the idea that glycosylation can play an important role in modulating macromolecular binding to the cell surface.

### Cell surface crowding is significant and varies across different cell types

Motivated by the unique insights obtained from our crowding sensors on human RBCs, we next applied them to quantify cell surface crowding of other mammalian cells, including tumor cells with upregulated glycosylation and sialylation (1). While our cholesterol-polymer sensors with adjustable size are useful for comparing different conditions for the same cell type, they cannot make comparisons across different cell types due to differences in lipid bilayer compositions and resulting incorporation level of the sensor. To overcome this challenge, we developed a new normalized crowding sensor that enables measurements across different cell types regardless of their size, shape, and lipid composition. The sensor consists of two parts – a cholesterol-biotin conjugate that spontaneously incorporates into the cell plasma membrane from solution (biotin anchor), and a fluorescent anti-biotin IgG antibody that measures cell surface crowding based on its binding affinity to the membrane-incorporated biotin (crowding sensor). The biotin anchor also contains a fluorescein (FITC) fluorophore via thymidine oligonucleotide linker to report its membrane incorporation level (5’-FITC-TTTTTT-biotin-TTT-cholesterol-3’) (Fig. 3A). The IgG size (~12 nm) (33,34) is similar to our 40k dextran sensor (~10 nm), and we confirmed that both sensors have similar sensitivity to surface crowding (Supplemental Fig. 3). Our new crowding sensor allows the simultaneous measurement of both the biotin surface density and anti-biotin antibody binding with single-cell resolution (Fig. 3B). The level of the surface-bound antibody concentration normalized by the biotin anchor signal, *θ*, as a function of the bulk antibody concentration provides a direct measurement of *K_D_* (Fig. 3C).

We next compared dissociation constants for different suspension and adherent mammalian cells, normalized by the value on bare beads, 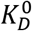(Fig. 3D). Interestingly, we find that all mammalian cells tested have surfaces that are significantly more crowded than any reconstituted surfaces evaluated in Fig. 1 (e.g. 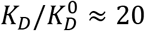 for SKBR breast epithelial tumor cells compared to 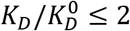 on bead surfaces with PEG3k, Fig. 1E). This highlights the importance of using native cell membranes to investigate physiological impacts of cell surface crowding.

Among the cells tested, the SKBR-3 breast cancer cells posed the largest crowding penalty against antibody binding. Like many epithelial tumor cells, SKBR-3 cells overexpress MUC1, a tall (200-500 nm), heavily-glycosylated mucin with 50-90% carbohydrate by mass (2,35–37). Shurer et al. demonstrated a direct correlation between MUC1 expression and cell membrane tubule formation, suggesting the MUC1 may be a major contributor to surface crowding (2). We speculate that while MUC1 can pose a steric barrier, shorter proteins also play a significant role in crowding near the membrane surface where the crowding sensor is located. In support of this hypothesis, we note that RBCs and HeLa cells have approximately the same 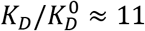, even though HeLa cells have a larger concentration of tall proteins, like MUC1, estimated to be at 10^4^ copies/HeLa cell on average (2,38). This, combined with the minimal crowding contribution of Fibcon-1 L we observed in Fig. 1F, suggests that our crowding sensor is most sensitive near the membrane surfaces, where bulky mucins likely contribute less to the values in Fig. 3D.

Interestingly, SKBR-3 displayed 2x more crowding compared to MCF10a cells, which are non-malignant breast epithelial cells commonly used to model normal breast epithelia behavior. This raises the question of whether reduced receptor binding near the cell surface due to upregulation of surface proteins plays a role in more malignant tumors. We did not find systematic correlations between our measurements and previously measured RNA expression level of the surface proteins, consistent with the notion that protein and RNA copy number are not well correlated for membrane proteins (28,39). Therefore, cell surface crowding, which varies significantly among different cells, cannot be predicted simply based on RNAseq databases.

### Cell surface crowding is altered by oncogenic mutations and surface protein overexpression

After observing the significant differences in cell surface crowding across different cell types, we asked whether mutations or changes in expression can alter crowding in a single cell type. To test if our crowding sensor could resolve such changes, we used lentivirus to generate HEK cell lines expressing or overexpressing surface proteins of different heights. Our measurements show that the surface crowding of cells expressing CD43, SIRPα, and E-cadherin increased crowding, while the crowding of Fib1L-expressing cell remains approximately the same (Fig. 4A). Interestingly, SIRPα expression increased crowding by approximately 2.5x compared to the wild type (Fig. 4B) and cell-cell crowding variability by ~25% (Fig. 4C), whereas CD43 expression increased crowding by less than a factor of 2x (Fig. 4B) and cell-cell crowding variability by ~ 40% compared to the wild type (Fig. 4C). We note that the crowding variability measured in the wild-type is ~60% larger than that in beads (Fig. 4C). This suggests that expression changes within a cell population can lead to both increased crowding and increased variability.

**Figure 4.**
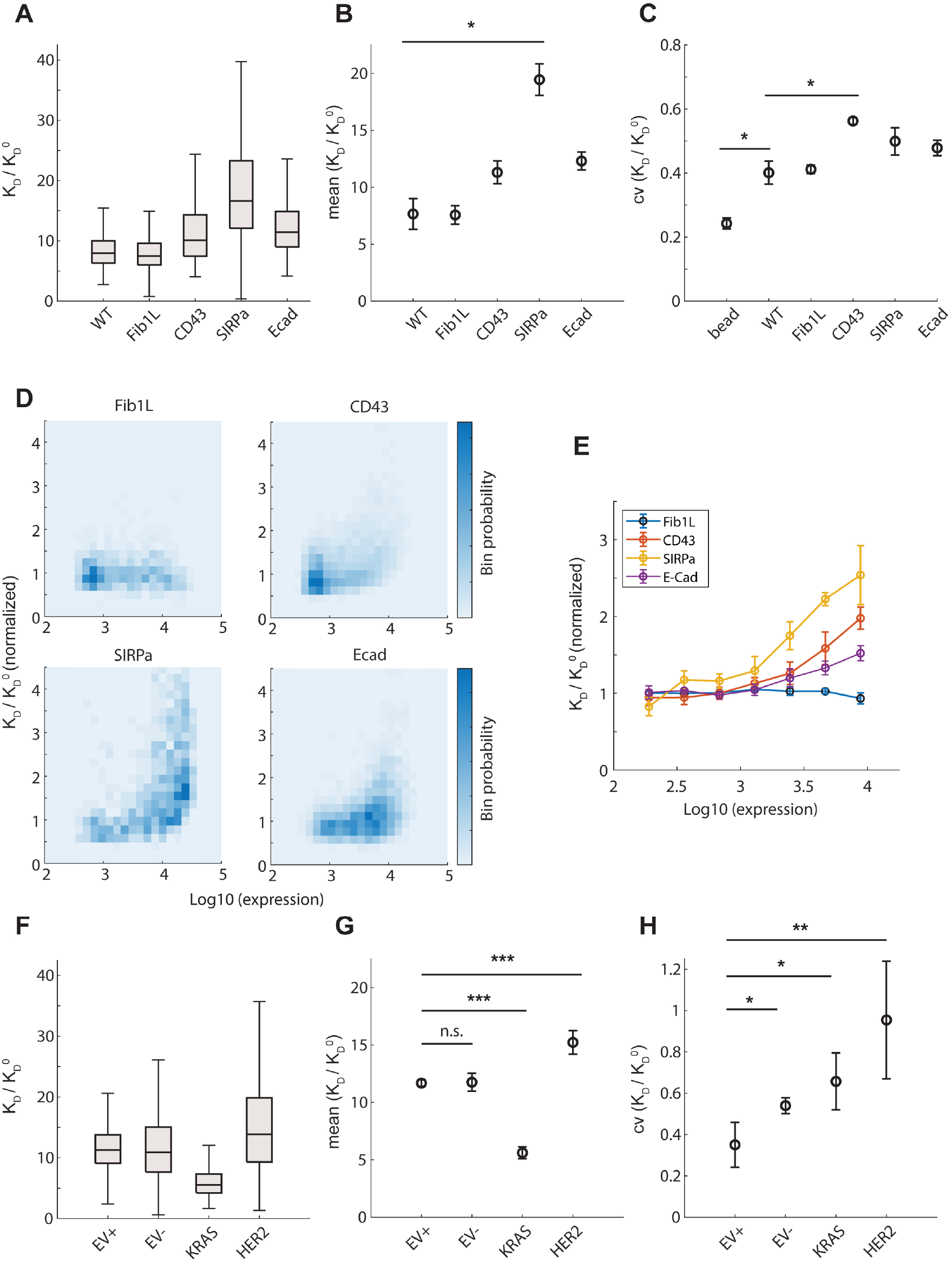
Mutations and surface protein overexpression alter cell surface crowding and variability. (A) Reduction in antibody binding affinity in cells overexpressing various proteins. The distribution of affinity change is represented as a box and whisker plot showing the median, lower and upper quartiles, and minimum and the maximum values ignoring outliers. N > 300 cells are measured in all conditions in a single run. (B and C) The mean and coefficient of variance (CV) of affinity change obtained from two replicate measurements. P-values are calculated based on a two-tailed *Student’s* t-test (* < 0.05). (D) Reduction in antibody binding affinity as a function of protein overexpression in the 2D bin scatter plot. (E) Reduction in antibody binding affinity as a function of protein overexpression. Error bars represent the standard deviation of the mean from three replicate runs. (F) Reduction in antibody binding affinity in normal MCF10a and MCF10a transformed with KRAS(G12V) and HER2. The distribution of affinity change is represented as a box and whisker plot showing the median, lower and upper quartiles, and minimum and the maximum values ignoring outliers. N > 300 cells are measured in all conditions in a single run. EV+: empty vector control supplemented with growth factors. EV-: empty vector control without growth factors (G and H) The mean and coefficient of variance (CV) of affinity change obtained from three replicate measurements. P-values are calculated based on a two-tailed *Student’s* t-test (* < 0.05, ** < 0.01, *** < 0.001).

To further examine the extent of surface crowding observed in different cell lines, we expressed the same surface proteins with a small, 12 residue Spot-tag fused at their N-terminus (16). We measured the change of crowding sensor binding as a function of protein expression level, which was inherently variable due to the lentivirus, by quantifying a fluorescent anti-spot nanobody (V_HH_) using a small ~2-4 nm antibody fragment. We found that crowding increased dramatically as a function of protein overexpression of CD43, SIRPα, and E-cadherin (Fig. 4D). We binned the cells by protein expression level and calculated crowding for each bin, which revealed that SIRPα increases the crowding the most for a given overexpression level, followed by CD43 and E-cadherin, while for Fib1L, little change was observed even over 2 orders of magnitude of expression (Fig. 4E). The extracellular domains of Fib 1L, SIRPα, CD43, and E-cadherin, are 89, 317, 204, and 543 amino acids, respectively, underscoring that surface crowding is determined not only by protein molecular mass but also contributions from properties such as surface charge and structure. Taken together, these results demonstrate that the change in a single protein’s expression can significantly alter the total cell surface crowding.

We next tested whether oncogenic transformation alters cell surface crowding using the breast epithelial cell line MCF10a expressing the common oncogenes HER2 and KRAS(G12V) (40). We first investigated the role of growth conditions. MCF10a requires growth factors for proliferation but can also maintain homeostasis in the absence of growth factors. We found that MCF10a cells exhibit similar levels of crowding in both in both proliferating and non-proliferating states (Fig. 4F and 4G). We measured the effect of the oncogenes and found that KRAS-expressing cells decreased crowding by 2x while HER2 cells show slightly increased crowding (Fig. 4G). Interestingly, cell surface crowding becomes significantly more variable for HER2 cells, with the top 25% reducing antibody binding affinity by more than 20x (Fig. 4F and 4H). Consistent with this, KRAS(G12V) and HER2 expression in MCF10a cells is known to significantly alter the surfaceome as well as glycosylation pattern (41). Furthermore, surface protease activity also changes in KRAS and HER2 cells, cleaving different surface protein groups (42). Our observations suggest that the molecular-level changes in transformed cells results in direct biophysical changes in their cell surface.

## Discussion

Traditional biochemical, genetic, and proteomics approaches excel at characterizing the molecular features of membrane proteins (28,39,43–45). However, these approaches cannot capture their multibody biophysical interactions on cell surfaces such as crowding. As a result, mechanistic understanding of the biophysical interactions that modulate the organization of the cell surface glycocalyx has been limited. In this work, we developed a simple experimental technique to quantify the impact of cell surface protein glycosylation, density, charge, stiffness, and other physical properties on macromolecular binding to the plasma membrane of live cells.

Our measurements provide a direct quantification of the steric energy barrier posed by a crowded cell surface. We found that these energies correspond to ~0.75-3 k_B_T for the case of IgG binding to buried receptors. These results provide a perspective on what cell surface “crowding” means and how it might be quantified. The free energy posed by the crowded cell surface arises from an osmotic pressure generated by the glycocalyx, given by Δ*U* ≈ *ΠV*^eff^, where Π is the osmotic pressure and *V*^eff^ is the effective volume occupied by the macromolecule within the glycocalyx (see Supplemental Information). The free energy may be interpreted as the mechanical work to displace a volume *V*^eff^ inside a crowded environment with pressure Π. All crowding contributions are captured by the osmotic pressure, including protein glycosylation, density, charge, stiffness, and other physical properties. In the Supplementary Information, we demonstrate using MD simulations that all sensor binding data collapse onto a universal curve described by a “cell surface equation of state” (Supplementary Fig. 1). We propose that the osmotic pressure is a universal metric that acts as a quantitative reporter of cell surface crowding, as opposed to other proxy metrics like protein molecular weight, surface charge, or number density.

The osmotic pressure measured proximal to the plasma membrane ranges from 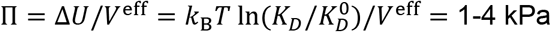, based on 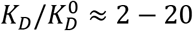 and an IgG with approximate size of 10 nm. These pressures are consistent with those generated by steric crowding interactions between synthetic polymer brushes on membranes, which are sufficient to bend the lipid bilayer (46–48). Interestingly, the surface osmotic pressures measured by our crowding sensors are larger than the stiffness of the cell cytoplasm (~100 Pa), the cell cortex (~1 kPa), and the thick glycocalyx of endothelial cells (100 to 500 Pa), as measured using atomic force microscopy with a large bead tip (49,50). We anticipate that the 1-4 kPa pressures are highly localized just above the membrane surface and that these pressures decay rapidly as a function of distance from the surface into the bulk fluid. Yet, large soluble ligands, antibodies, viruses, and receptors from opposing cell surfaces that need to bind close to the membrane surface will experience large osmotic pressures opposing binding due to crowding of the glycocalyx. Indeed, we found that the surface crowding can reduce the binding isotherm of an IgG antibody to a cell surface by as much as 20x.

We end by noting several areas for future investigation. First, we focused on crowding measurements at a fixed distance just above the membrane surface. Since the glycocalyx is a fully 3D structure, we anticipate that the surface crowding is height dependent and a crowding measurement as a function of distance from the membrane may provide further insight into cell surface organization. Second, our sensors may be used to study temporal changes in surface protein expression during different stages of the cell cycle, tumor progression, cell senescence, and cell differentiation. Third, while we focused on the extracellular side of the cell membrane, the cytoskeleton on the interior side plays a key role in membrane protein organization, and the interplay between cortical actin organization and glycocalyx crowding mediated by actin-binding proteins merits further study. Lastly, despite the advances in quantitative transcriptomics and proteomics (28,39,43–45), it remains unclear whether and how protein copy number on a cell surface is regulated under physiological conditions or whether surface crowding is simply an unregulated outcome of protein expression, glycosylation, and trafficking. Our new crowding sensors may be used to address these and other topics related to the biophysical properties and collective function of the cell surfaceome.

## Materials and Methods

### Materials

Purified human Glycophorin A (CD235a) extracellular domain with a C-terminal 6x-His tag (catalog number: 16018-H08H) was purchased from Sino Biological US Inc.

Alexa Fluor 647 labeled anti-biotin (BK-1/39; catalog number: sc-53179 AF647) was purchased from Santa Cruz Biotechnology. Monoclonal antibodies against human Glycophorin AB (CD235ab) (HIR2; catalog number: 306602) and human CD47 (CC2C6; catalog number: 323102) were purchased from BioLegend.

BD Microtainer Contact Activated Lancet, 30 G x 1.5 mm, low flow, purple (catalog number: 366592) was purchased from BD Biosciences.

Lectins Sambucus Nigra Lectin (SNA; catalog number: L-1300), Fluorescein labeled Agaricus bisporus lectin (ABL; catalog number: FL-1421), Fluorescein labeled Erythrina cristagalli lectin (ECL; catalog number: FL-1141), and Wheat Germ Agglutinin (WGA; catalog number: L-1020-10) were purchased from Vector Laboratories.

1,2-dioleoyl-sn-glycero-3-(N-(5-amino-1-carboxypentyl)iminodiacetic acid)succinyl (DGS-Ni-NTA; catalog number: 709404), 1,2-dioleoyl-sn-glycero-3-phosphocholine (DOPC; catalog number: 850375), 1,2-dioleoyl-sn-glycero-3-phospho-L-serine (DOPS; catalog number: 840035), 1,2-dioleoyl-sn-glycero-3-phosphoethanolamine-N-(cap biotinyl) (Biotinyl Cap PE; catalog number: 870273), 1,2-dioleoyl-sn-glycero-3-phosphoethanolamine-N-methoxy(polyethylene glycol)-2000 (PEG2k PE; catalog number: 880130), 1,2-dioleoyl-sn-glycero-3-phosphoethanolamine-N-methoxy(polyethylene glycol)-3000 (PEG3k PE; catalog number: 880330) were purchased from Avanti polar lipids.

Silica microspheres (4.07μm; catalog code: SS05002; lot number: 12602 and 6.84μm; catalog code: SS06N; lot number: 4907) were purchased from Bangs laboratories. Cholesterol NHS (catalog number: CSL02) was purchased from Nanocs Inc. Cholesterol-PEG-Amine, MW 1k (catalog number: PLS-9961) was purchased from Creative PEGWorks. Dextran, Amino, 10,000 MW (10k dextran; catalog number: D1860), Dextran, Amino, 40,000 MW (40k dextran; catalog number: D1861), Alexa Fluor 647 NHS Ester (Succinimidyl Ester) (NHS-AF647; catalog number: A20006), Alexa Fluor 555 NHS Ester (Succinimidyl Ester) (NHS-AF555; catalog number: A20009), Alexa Fluor 488 NHS Ester (Succinimidyl Ester) (NHS-AF488; catalog number: A20000), BODIPY FL NHS Ester (Succinimidyl Ester) (NHS-BODIPY; catalog number: D2184) were purchased from Invitrogen.

NHS-Fluorescein (5/6-carboxyfluorescein succinimidyl ester) (NHS-FITC; catalog number: 46409), DyLight 650-4xPEG NHS Ester (catalog number: 62274), EZ-Link Hydrazide-Biotin (hydrazide-biotin; catalog number: 21339), EDC (1-ethyl-3-(3-dimethylaminopropyl)carbodiimide hydrochloride) (catalog number: 77149), Zeba Spin Desalting Column, 7K MWCO (catalog number: 89882) were purchased from Thermo Scientific.

Proteinase K from Tritirachium album (catalog number: P2308) and sialidase from Clostridium perfringens (C. welchii) (catalog number: N2876) were purchased from Sigma.

MATLAB educational license was obtained from MathWorks Inc. UC Berkeley’s BRC High Performance Computing Savio cluster with NVIDIA K80 GPU were used for molecular simulations. Attune NxT Acoustic Focusing Cytometer (ThermoFisher Scientific) was used for all of the flow cytometer experiments.

### Protein purification

Multi-FN3-domain proteins containing C-terminal 10x His-tag and N-terminal ybbR-tags were purified as described previously (20). Briefly, FN3 proteins were expressed in Rosetta DE3 cells (EMD Millipore), lysed by sonication, and purified over a His-Trap HP column (GE Healthcare). The proteins were gel-filtered and their size was confirmed via a Superdex 200 column on an AKTA Pure system (GE Healthcare).

### Microscopy

All imaging was carried out on an inverted Nikon Eclipse Ti microscope (Nikon Instruments) equipped with a Yokogawa CSU-X spinning disk using an oil-immersion objective (Apo TIRF 60x and 100x, numerical aperture (NA) 1.49, oil). Three solid state lasers were used for excitation: 488nm, 561nm, and 640nm (ILE-400 multi-mode fiber with BCU, Andor Technologies). The laser power at the sample plane was less than 1.5mW for all three channels. Fluorescent light was spectrally filtered with emission filters (535/40m, 610/75, and 665LP, Chroma Technology) and imaged on a sCMOS camera (Zyla 4.2, Andor Technologies). For CSOP measurements, Z-stack was acquired using a piezo z-stage (npoint).

### Synthesis of crowding sensors

Cholesterol NHS (Nanocs) was dissolved in 1:2 ratio of ethanol and DMSO. Amino 10k dextran and 40k dextran (Invitrogen) and Cholesterol-PEG-amine (Creative PEGWorks, Inc; 966 g/mol MW) were dissolved in DMSO. Equimolar ratio of NHS-dye (choice of AF488, AF555, AF647, or BODIPY), and cholesterol NHS were mixed at a 10:1 molar ratio with cholesterol-PEG-amine, 5:1 molar ratio with 10k dextran amino, 20:1 molar ratio with 40k dextran amino, and left overnight at 50C. Control probes without cholesterol conjugation were mixed without cholesterol NHS. To remove unreacted NHS reactants, the reaction mixture was processed through a Zeba Spin Desalting Column, 7K MWCO (Thermo Scientific). Labeling ratio of cholesterol and dye were recorded using a NanoDrop 2000c spectrophotometer (Thermo Scientific).

Small aliquots were stored in −80C and used within a few months. Thawed sensors were used within the same day. The labeling ratio of the small sensors is 1 cholesterol/dye, 10k sensor is 3-5 cholesterol/AF555, and 40k sensor is 7-10 cholesterol/AF647. The approximate hydrodynamic diameters of the Chol-PEG-dye, 10k, and 40k sensors are ~2, 4, and 10 nm, respectively (19).

### Preparation of supported lipid bilayer on glass beads

Lipid-coated glass beads were created by coating glass micro-beads with a fluid supported lipid bilayer (SLB). Small unilamellar vesicles (SUVs) were prepared by rehydrating a lipid sheet composed of DOPC and other phospholipids with pure deionized water. For SLBs involving the attachment of His-tagged purified proteins, the lipids were mixed with 7.5% of DGS-Ni-NTA. For SLBs involving the binding of anti-biotin antibody, the lipids were mixed with 7.5% of DGS-Ni-NTA, 1% biotinyl cap PE, and 1% of PEG2k PE or PEG3k PE. For all other SLBs, the lipids were mixed with 5% DOPS, and up to 1% of PEG3k PE.

After rehydrating for 30 min, the solution was vigorously vortexed, sonicated at low-power (20% power) using a tip-sonicator, and finally filtered through a 0.2 mm PTFE filter (Millipore). Stock solutions of SUVs were stored at 4C and were used within 48 hours to avoid phospholipid oxidization.

4.07 μm and 6.46 μm glass micro-bead (Bangs labs) slurry (10% solids) were cleaned using a 3:2 mixture of H2SO4:H2O2 (Piranha) for 30 minutes in a bath sonicator, and were spun down at 1000g and washed 3 times before being resuspended in pure water. Clean beads were stored in water at room temperature and used within 48 hours. To form SLBs, 45 μL of SUV solution was added with 5 μL of 10x MOPS buffer (500mM MOPS pH 7.4, 1M NaCl) and 10μL of clean bead suspension, and mixed gently. The bead/SUV mixture was incubated for 15 minutes at room temperature while allowing the beads to sediment to the bottom of the tube. Beads were washed 5 times with HEPES buffer (50mM HEPES pH 7.4, 100mM NaCl) by gently adding/removing the buffer without resuspending the glass beads into solution.

We verified the fluidity of the lipid bilayer by imaging beads on a glass coverslip with a spinning-disk confocal microscope (Nikon) at 60x magnification and high laser intensity, where diffusion of labeled lipids was visible after photo-bleaching a small region.

For experiments involving poly His-tagged purified Fibcon and GYPA proteins, 200nM of protein was added into the lipid-coated bead solution and incubated at room temperature for 20 minutes. Beads were washed 3 times with HEPES buffer by gently adding/removing the buffer without resuspending the glass beads into solution. For sialidase-treated GYPA beads, the beads were further treated with 200 mUn/mL of sialidase for 30 minutes at 37C. For pro-K treated beads, the beads were further treated with 0.05 mg/mL of pro-K for 20 minutes at 37C. Beads were washed 3 times with HEPES buffer.

### Preparation of red blood cells

BD Microtainer contact-activated lancet was used to withdraw blood from volunteers. A small amount (10 μL) of blood was washed in phosphate buffered saline (PBS) with 2 mM ethylenediaminetetraacetic acid (EDTA) to remove soluble proteins and plasma from whole blood. Red blood cells were stored in PBS solution at 4C and used within a day. All procedures followed a UC Berkeley IRB approved protocol (CPHS Protocol Number: 2019-08-12454).

To cleave sialic acid from the surface of RBCs, sialidase was used at 50 mUn/mL in PBS at 37C for 2 hours. No protease activity was found in our stock NA, as shown in the sodium dodecyl sulfate polyacrylamide gel electrophoresis (SDS-PAGE) protein gel (see SI). To digest the cell surface proteins of RBCs, pro-K was added at 0.05 mg/mL in PBS at 37C for 1 hour. To remove negative charges on carboxylic acid groups on red blood cell surfaces, 1mM hydrazide-biotin and 30mM EDC were mixed with untreated cells in PBS at room temperature for 6 hours. The cell surface was visualized with confocal microscopy using fluorescently-labeled streptavidin that binds to biotin (see SI).

RBC measurements were conducted using the synthetic dextran-based sensors. Small Chol-13xPEG-488 sensor was varied between 0-10 nM, the medium 10k-555 sensor was varied between 0-15nM, and the large 40k-647 sensor was varied between 0-5 nM.

### Flow cytometry

An Attune NxT Acoustic Focusing Cytometer (ThermoFisher Scientific) was used for all flow cytometry experiments. For RBC measurements, we added ≈ 10^6^ RBC into 1 mL of PBS containing appropriate sensors (typically ≈ 30 *μL* of stock RBC, containing 10 *μL* of collected blood in 400 *μL* of PBS). Sensors were incorporated into cells by incubating with the appropriate concentration, generally between 20-40nM.

Since the synthetic crowding sensors and the monoclonal antibodies are fluorescently labeled, the intensity units from the flow cytometer directly reports on the bound sensor concentration. The fractional surface coverage is defined as *θ* = *C/C^max^*, where *C* is the surface concentration of the sensor on a given sample averaged across ~20000 events. This fractional surface coverage is plotted as function of bulk concentration of sensor, and the slope at small bulk sensor concentrations directly gives the dissociation constant.

### Cell Surface Optical Profilometry (CSOP) height measurements on RBCs

PBS solution containing unconjugated WGA was incubated into an 8-well chambered cover glass (Cellvis; catalog number: C8-1.5H-N) to coat the glass with WGA. This helps to immobilize the RBCs when added to the chamber. The chambers were washed and filled with 0.25x PBS in MQ water to swell the RBC into a bloated sphere. Our Chol-13xPEG-488 sensors were added (83nM) as a membrane reporter. Untreated and sialidase-treated RBCs were added to separate wells. We acquired approximately 15 slices of images straddling the bead equator with a 100 nm step size. Each z-stack image contained up to ten beads per field-of-view. It is critical to locate the equator in both channels to ensure that accurate offsets are calculated (16). More than 100 RBCs were acquired and processed in our custom MATLAB script. Cells with visible defects, including non-spherical shape, membrane tubule and/or bud formation were removed from analysis. In addition, chromatic aberration and other optical offsets were subtracted from our signal by taking a baseline calibration measurement on RBC using our membrane sensor pairs Chol-13xPEG-488 at 83nM and Chol-13xPEG-647 at 125nM. The height of the cell surface proteins is obtained by quantifying the difference between the fluorescently-labeled height and the aberration offset, 〈*h*〉 = 〈*h*_measured_〉 – 〈*h*_offset_〉.

The N-terminal alpha-amines of red blood cells were labeled with DyLight 650-4xPEG NHS at 100 μM for 15 minutes at room temperature in PBS at pH 6.5 (with citric acid). NHS reaction at low pH facilitates preferential labeling of the N-terminus (average *pK_a_* ≈ 5 - 7) as opposed to other aliphatic amines on the protein (average *pK_a_* ≈ 10.5) and phosphoethanolamine lipids (*pK_a_* > 10) (51). The reaction mixture was washed with PBS to remove excess NHS reagent. Proteinase-K treatment of these cells showed that the majority of the label was removed from the RBC surface, verifying that the NHS reaction predominantly labels the proteins and not the lipids. For sialidase treatment, the cells were treated with 50 mUn/mL for 1 hour at 37C to digest sialic acids from the cell surfaces prior to CSOP measurement. No divalent cations (including calcium and magnesium) were added in the measurements, which could result in unwanted crosslinking between negative charges in the glycocalyx.

In addition to N-terminal labeling of RBC surface, we also used lectins as a readout of cell surface height. For height measurements based on lectins, fluorescein labeled ABL, and ECL were used. ABL and Jacalin bind to galactosyl (beta-1,3) N-acetylgalactosamine (also called the Thomsen-Friedenreich antigen, or T disaccharide), which are heavily expressed on glycophorin proteins on the RBC surface. Unlike peanut agglutinin, which does not bind sialylated T antigen, ABL bind either sialylated or asialylated forms, which makes them good markers for height measurements. Although we notice a 30-50% increase in the binding of these lectins to sialidase-treated RBCs, we still observe a strong signal on the fully sialylated surface and assume that the spatial distribution of bound lectins does not change significantly between untreated and sialidase-treated RBCs. ECL binds to the disaccharide Gal (*β*1 - 4) GlcNAc, called N-acetyllactosamine (LacNAc). Although ELC does not bind to LacNAc terminated with sialic acids, we still observed strong binding on red cells for all conditions. The single N-glycosylation on the Band3 protein contains LacNAc, which is abundant on the RBC surface. The single complex N-glycan on Band3 is heterogeneous in size on different molecules, varied by the repeat of poly-LacNAc units (Gal- *β*1 → 4 GlcNAc *β*1 → 3). The end of the oligosaccharide can be linked to sialic acid, fucose, or exposed.

### Coarse-grained molecular dynamics (MD) simulations of protein polymers and sensors

To construct a molecular model of macromolecular transport across cell surface proteins and glycocalyx, we performed coarse-grained Molecular Dynamics (MD) simulations of particle transport within semi-flexible polymers diffusing on 2D surfaces. See Supplementary Information for a detailed explanation of our simulations.

Briefly, we model surface proteins using a Kremer-Grest bead-spring model (52), with each bead representing a structured protein domain or a coarse-grained unit of an intrinsically disordered domain. Because the size of each protein domain is large compared to the surrounding solvent molecules, the solvent is coarse-grained out and its dynamics are not explicitly evolved. In other words, the protein chain experiences a hydrodynamic drag and Brownian motion from the continuous solvent. In this work, the membrane does not deform nor fluctuate in the out-of-plane dimension, although these effects may be included.

Simulations were performed using a GPU-enabled HOOMD-blue molecular dynamics package (53). A dilute concentration of soluble particles with different sizes were added to model the dynamics of sensor binding to the cell surface. The diameters of the Chol-PEG, 10k, and 40k sensors were modeled with spheres of diameter 3, 5, and 10 nm, respectively. The sensor particles have an attractive potential to the membrane via a shortrange, attractive harmonic potential *U*_att_ = *k*(*z* - *r*_0_)^2^/2 with stiffness *k*. All other particle pairs experience a short-ranged repulsive Weeks-Chandler-Andersen (WCA) potential. Charges were included into the model by adding a screened Coulomb (Yukawa) potential between pairs of particles, with the Debye length as an additional parameter.

We imposed periodic boundary conditions in the *x* and *y* directions, and no-flux hard walls at *z* = 0 and *z* = *L_z_* to prevent any particles from passing through. Initial configurations were generated by placing the particles in lattice locations, and sufficient time steps are run to reach equilibrium. We verified that the chosen time steps (Δ*t* = 10^-6^ – 10^-4^) are sufficiently small to capture the relevant dynamics.

### Mammalian cell culture and preparation

HEK293T cells were obtained from UCSF Cell Culture Facility and grown in DMEM (Life Technologies) supplemented with 10% heat-inactivated FBS (Life Technologies) and 1% Pen-Strep (Life Technologies), at 37°C, 5% CO2.

SK-BR-3 cells were purchased from ATCC and grown in McCoy’s 5A (ATCC) supplemented with 10% FBS and 1% Pen-Strep.

Jurkat and L1210 cells were purchased from ATCC and is grown in RPMI (Life Technologies) supplemented with 10% FBS, 1% Pen-Strep.

Hela cells were obtained from UC Berkeley Cell Culture Facility and grown in DMEM (Life Technologies) supplemented with 10% heat-inactivated FBS (Life Technologies) and 1% Pen-Strep (Life Technologies), at 37°C, 5% CO2.

MCF10a-derived cell lines expressing oncogenes were a kind gift from Kevin Leung and James Wells at UCSF (41).

For depletion experiments, cells were grown in DMEM supplemented with 5% horse serum, 0.5mg/mL hydrocortisone, 100ng/mL cholera toxin, and 1% Pen-Strep. For proliferation experiments, media was further supplemented with 20ng/mL Epidermal Growth Factor and 10μg/mL insulin. Cells were passaged by treatment with 0.05% trypsin, and for depletion experiments, by treatment with versene.

Cells were passaged every 2-3 days. One day prior to flow cytometry experiments, care was taken to seed adherent cells at dilute to intermediate concentrations to prevent the formation of multi-cell clusters.

For suspension cells (red blood cells, L1210, Jurkat), cells were washed by centrifuging at 150xg for 1 minute and resuspending in PBS to remove media. Adherent cells were carefully scraped from the plastic substrate using a cell scraper and then washed.

To prevent internalization of the macromolecules into the cell interior, cells were then incubated on ice for 10-15 minutes, and all subsequent steps were performed on ice. Then, 10-100nM of the FITC-Biotin-Cholesterol synthetic antigen sensor was incubated with the cells for 20 minutes. The final isotherm results for the anti-biotin antibody are insensitive to the precise amounts of FITC on the cell surface, as long as the total FITC concentration is low (less than ~200 / μm^2), as evidenced by the fixed slope of the 488 vs 640 intensity in the raw flow cytometry data (Fig. 3B). The cells were washed thoroughly (3x) in PBS in a centrifuge set to 4C. Finally, the cells were added into tubes prepared with several different AF647-labeled anti-biotin monoclonal antibody concentration (0 −0.5μg/mL). After incubation on ice for 40 minutes, the samples were measured on the flow cytometer.

### Lentiviral preparation and cell line generation

Lentivirus was produced by transfecting the transfer plasmids, pCMV-dR8.91, and pMD2.G (1.5μg, 1.33μg, and 0.167μg per 35mm well) into 293T cells grown to approximately 80% confluency using Mirus TransIT-293 Transfection Reagent (Promega) per manufacturer’s protocol. After 60-72 hours, supernatant containing viral particles was harvested and filtered with a 0.45 μm filter. Supernatant was immediately used for transduction or aliquoted and stored at −80°C. Cells were seeded at 20% confluency in 35mm dishes and 0.5-1mL of filtered viral supernatant was added to the cells. Media containing virus was replaced with fresh growth medium 24 hr post-infection. Infected cells were imaged to assess transduction efficiency and then used in flow cytometry assays as described above.

## Acknowledgments

The authors would like to thank Kevin Leung and James Wells at UCSF for a kind gift of MCF10a-derived cell lines. S.C.T. is supported by the Packard Fellowship in Science and Engineering and the National Science Foundation (Grant No. 2150686). D.A.F. is supported by NIH R01 GM134137 (DAF), the NSF Center for Cellular Construction (DBI- 1548297), and the Miller Institute for Basic Research. D.A.F. is a Chan Zuckerberg Biohub Investigator.

## SUPPLEMENTAL FIGURES

**Supplemental S1.**
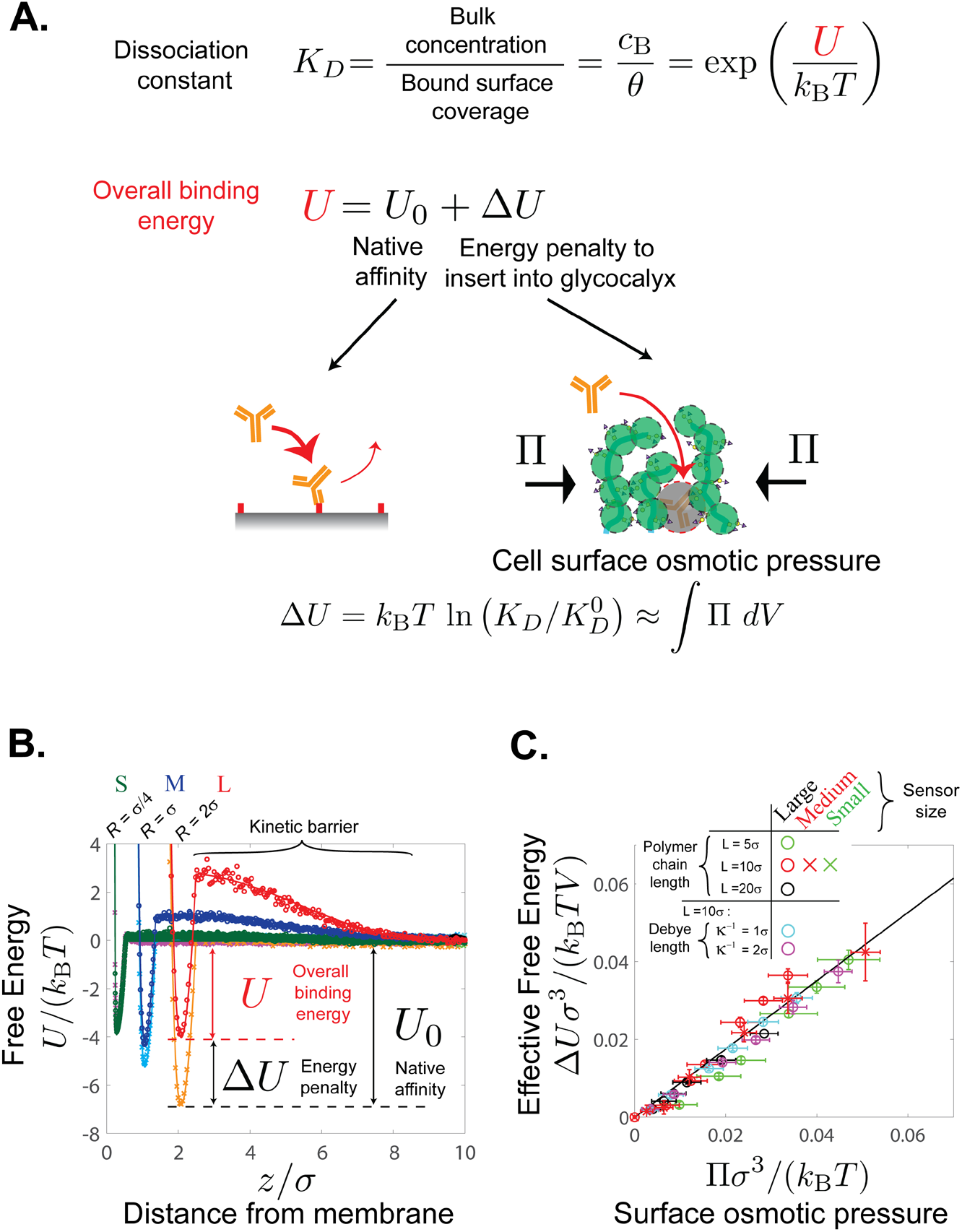
Macromolecular crowding on membrane surfaces is described by a surface osmotic pressure. (A) The dissociation constant of a surface-binding macromolecule is given by its binding energy, *U*, which is a sum of the intrinsic affinity, *U*_0_, and the penalty due to crowding, Δ*U*. The energy penalty posed by surface crowding is directly related to the change in normalized binding affinity, 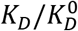, and the osmotic pressure of the surface, Π. (B) Coarse-grained molecular dynamics (MD) simulations were used to calculate the binding energy curves for spherical sensors of sizes *σ_s_* = 0.25*σ* (small), *σ* (medium), and 2*σ* (large), where *σ* is the coarse-grained effective diameter of a glycocalyx polymer chain. The depth of the minimum gives the effective binding energy at equilibrium. The surface contains polymers with contour length 10σ at a concentration of ~ 10000/*μm*^2^. (C) The energy penalty across various sensor sizes, surface polymer lengths, and surface polymer charges all collapse onto a unifying scaling line when the data are plotted against the osmotic pressure generated by the surface polymers, Π. The multi-domain proteins on the surface have varying lengths and are either charged (cyan and magenta circles) or neutral (all other symbols). Electrostatic interactions among the polymers are modeled by a Yukawa potential with different Debye lengths, *κ*^-1^. The energy penalty Δ*U* is normalized by the effective volume of the sensor immersed inside the crowded surface, *V*, and Π is calculated directly via the Irving-Kirkwood virial stress tensor. Error bar indicates standard deviations of ensembles over all time and particles. The solid line is given by analytical theory and is not a fit (see Supplementary Information).

**Supplemental S2.**
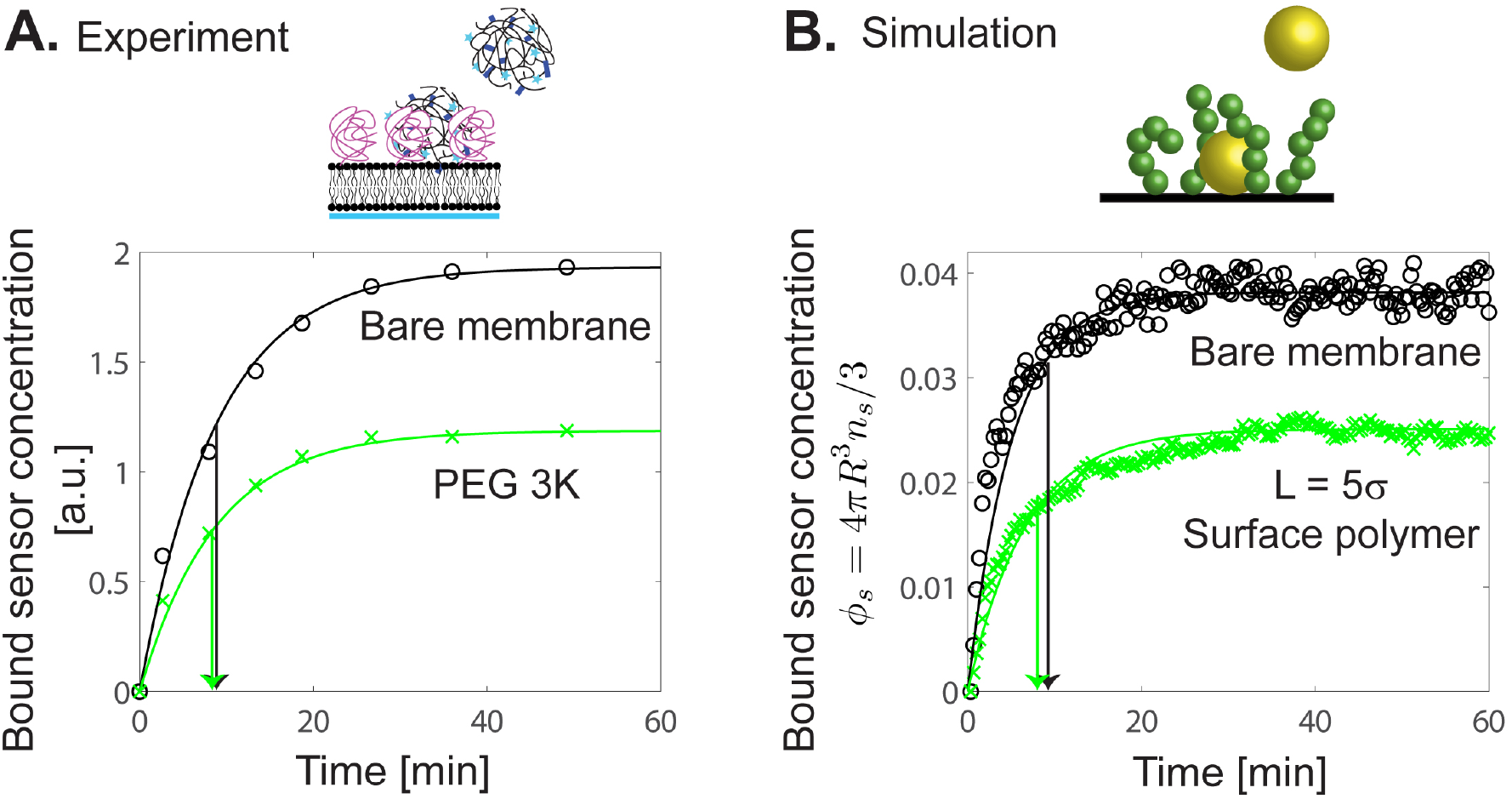
Sensor binding on crowded membrane surfaces is not transport limited. (A) The bound sensor concentration as a function of time on bare membrane beads (black symbols) and crowded PEG 3K surfaces (green symbols) saturates with the same time constant. Solid curves are a fit to a first-order kinetic rate law, *C/C*_∞_ = 1 - exp(- t/ *τ*). The characteristic decay time towards equilibrium is *τ* = 8.86 min for bare membranes and *τ* = 8.74 min for crowded PEG 3K surfaces. (B) MD simulations of sensor binding on bare surfaces (black symbols) and crowded surfaces (green symbols), where *ϕ_s_* = 4*πR*^3^*n_s_*/3 is the volume fraction of bound sensors at the surface. The characteristic decay time towards equilibrium is *τ* = 9.8 min for bare membranes and *τ* = 9.0 min for crowded polymer surfaces.

**Supplemental S3.**
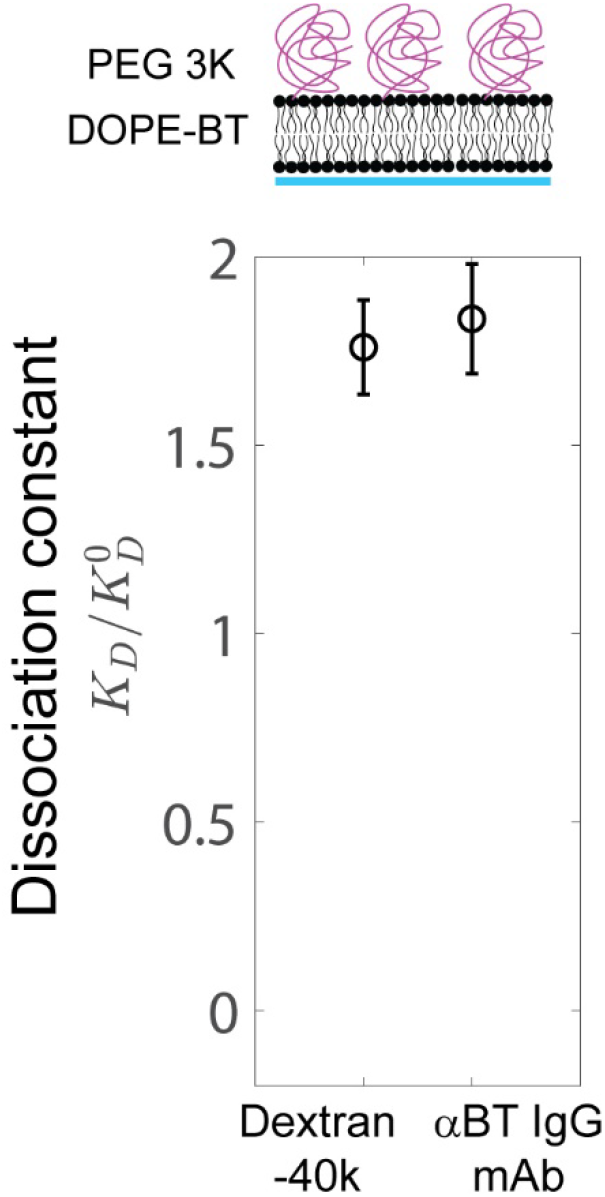
The crowding penalties reported by the large dextran-40k sensor and anti-biotin IgG antibodies are quantitatively similar on crowded reconstituted surfaces, consistent with their similar size (~10 nm). We confirmed that our alternative measurement using antibodies provides a proficient readout of crowding on live cells regardless of their size, shape, and lipid composition.

