## Supplementary Information for "Engineered molecular sensors of cell surface crowding"

### Chapter I. Dynamics of macromolecular binding on crowded surfaces

The goal of this chapter is to develop a theory of macromolecular adsorption to identify a quantitative metric of surface crowding. We aim to obtain a direct relationship between surface binding of soluble macromolecules and the crowding state of the surface, including the effects of variable glycoprotein length, polydispersity, stiffness, density, charge, and all other interactions. We propose that the osmotic pressure is a universal metric that acts as a quantitative reporter of cell surface crowding, as opposed to other proxy metrics like protein molecular weight or number density.

#### 1. Momentum balance on crowded cell membrane surfaces

The Cauchy momentum balance over a control volume just above the cell surface is given by

$$\rho \frac{D\boldsymbol{u}}{Dt} + \nabla \cdot \boldsymbol{\sigma} + \boldsymbol{b} = \mathbf{0}, \tag{1.1}$$

where $D/Dt$ is a material derivative, $\boldsymbol{u}$ is the suspension average velocity, $\boldsymbol{\sigma}$ is the total stress of the suspension, and $\boldsymbol{b}$ is an external force. Incompressibility further requires that $\nabla \cdot \boldsymbol{u} = 0$. The total suspension stress, which include both the fluid and the surface polymers, is given by

$$\boldsymbol{\sigma} = -p_f \boldsymbol{I} + 2\eta_0(1 + 5\phi/2)\boldsymbol{E} + \boldsymbol{\sigma}^{(P)}, \tag{1.2}$$

where $p_f$ is the fluid pressure, $\eta_0$ is the viscosity of the continuous Newtonian solvent, $\boldsymbol{E}$ is the rate-of-strain tensor, $\phi$ is the volume fraction of particles ($= 4\pi a^3 n/3$ for spheres), $n$ is the number density, $5\phi/2$ is the Einstein shear viscosity correction that is present for all suspensions, and the particle contribution to the stress $\boldsymbol{\sigma}^{(P)} = -nk_BT\, \boldsymbol{I} + \boldsymbol{\sigma}^P$, where $\boldsymbol{\sigma}^P = -n\langle \boldsymbol{x}_{ij}\boldsymbol{F}_{ij}\rangle$ is the virial stress contribution from interparticle interactions.

At low Reynolds numbers and in the absence of external forces, the momentum balance becomes

$$\nabla \cdot \boldsymbol{\sigma} = \mathbf{0}. \tag{1.3}$$

Assuming that there are no shear stresses and that the system is translationally invariant (i.e., independent of in-plane coordinates $x$ and $y$), we obtain $\partial\sigma_{zz}/\partial z = 0$.

This reveals that the stress is constant everywhere along the z-direction (normal to the membrane surface). Furthermore, in the z-direction, the steric repulsion due to polymer interactions exactly cancels the restoring force due to chain stretching, and $\sigma_{zz}^{(P)} = 0$. We therefore conclude that the fluid pressure $p_f$ is uniform and constant everywhere in the system.

In the x- and y-directions (tangent to membrane surface), the particle stress can take on any value, $\sigma_{xx}^{(P)} \neq 0$ and $\sigma_{yy}^{(P)} \neq 0$. We therefore have an anisotropic particle stress tensor, $\boldsymbol{\sigma}^{(P)} = \sigma_{xx}^{(P)}\boldsymbol{e}_x\boldsymbol{e}_x + \sigma_{yy}^{(P)}\boldsymbol{e}_y\boldsymbol{e}_y + \sigma_{zz}^{(P)}\boldsymbol{e}_z\boldsymbol{e}_z$, with $\sigma_{zz}^{(P)} = 0$. The mechanical (osmotic) pressure is defined as $\Pi = -\mathrm{tr}\,\boldsymbol{\sigma}/3$, and we define the in-plane pressure of cell surfaces as

$$\Pi = -\left(\sigma_{xx}^{(P)} + \sigma_{yy}^{(P)}\right)/2\,. \tag{1.4}$$

This is the in-plane pressure that we reported in Supplementary Fig. 1 and in this document from both experiments and simulations.

#### 2. Equation of state (EOS) of crowded membrane surfaces

Modeling the cell surface proteins and glycans as coarse-grained polymers, we can obtain an equation of state relating the osmotic pressure of the cell surface suspension as a function of its material properties, including density, contour length, persistence length, and electrostatic charge.

The equation of state for hard-sphere colloidal suspensions, including the Carnahan-Starling EOS, is used commonly to model macromolecular crowding on the membrane surface and inside the cytoplasm (1-4). Here, we use our data from molecular dynamics (MD) simulations to determine the equation of state for cell surfaces (see Chap. II for details of MD simulations). The EOS may be represented as a virial expansion(5):

$$\Pi = \frac{k_B T}{V_p}\left(\frac{\phi}{N_R} + B_2\phi^2 + B_3\phi^3 + \cdots\right), \quad (1.5)$$

where $V_p = \pi\sigma^3/6$ is the volume of a monomer, $N_R = L/\sigma$ is the degree of polymerization, $B_2$ and $B_3$ are the second and third virial coefficients and describe two-body and three-body interactions, respectively. For Weeks-Chandler-Andersen (WCA) potentials between all particles, $B_2 > 0$ and $B_3 > 0$, and the pressure increases beyond the ideal-gas value.

As shown in Fig. 1.1, we find that truncation at the three-body level gives a proficient agreement over the concentrations that we tested in MD. For charge-neutral polymer surfaces, $B_2 = 0.40$, $B_3 = 7.93$. For charged polymers with Debye length $\kappa^{-1} = 1\sigma$, we obtain $B_2 = 2.22$, $B_3 = 10.81$; for $\kappa^{-1} = 2\sigma$, we obtain $B_2 = 2.68$, $B_3 = 29.67$.

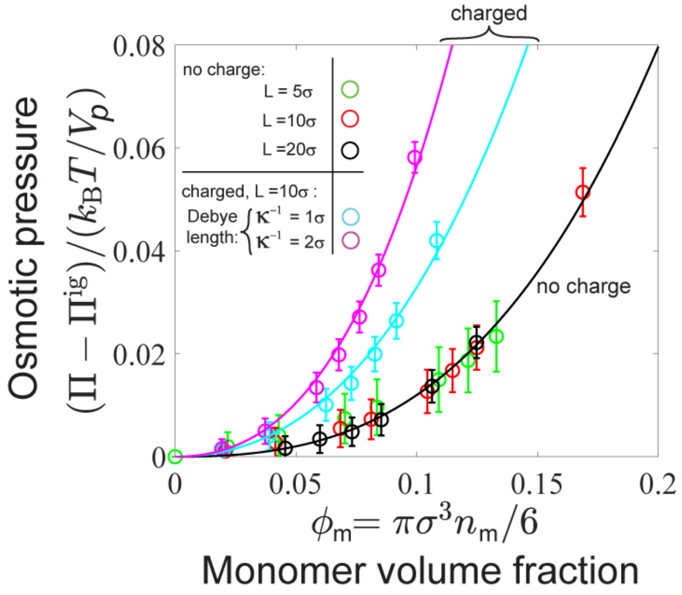


**Figure 1.1:** Coarse-grained molecular dynamics (MD) simulations were used to compute the osmotic pressure generated by polymers tethered to a membrane surface. The osmotic pressure is defined by Eqs. 1.4 and 1.5, and the ideal-gas pressure has been subtracted in this plot, $\Pi^{ig} = k_B T\phi/(V_p N_R)$, where $V_p$ is the volume of a monomer and $N_R = L/\sigma$ is the degree of polymerization. The symbols are simulation data, and the solid curves are Eq. 1.5.

#### 3. Analytical theory for brush inclusion penalty

Halerpin et al. (6-8) have conducted theoretical analyses of soluble protein adsorption onto chemically grafted polymer brush surfaces. They distinguish between an insertion versus compressive mechanism of protein adsorption, depending on the size of the protein compared to the mesh size of the polymer brush. One key difference between chemically grafted polymer brushes and the cell surface glycoprotein brush is that a significant fraction of transmembrane proteins is mobile and can translate in 2D along the lipid bilayer surface. Therefore, a large macromolecule may adsorb onto the membrane via the insertion mechanism because it is energetically more favorable to exclude polymers from the interface as opposed to compressing the brush.

To obtain the free energy of inserting a large macromolecule into the cell surface glycocalyx, we follow the theory by Louis et al (9). The inclusion energy of a colloid in a polymer suspension is given by

$$\Delta \mathrm{U} = \Pi(\phi)\mathrm{V_s} + \gamma(\phi)\mathrm{A_s}, \tag{1.6}$$

where $\Pi(\phi)$ is the osmotic pressure, $\gamma(\phi)$ is the interfacial tension, $\mathrm{V_s}$ is the volume of the macromolecule (= $4\pi R^3/3$ for spheres), and $\mathrm{A_s}$ is the surface area of the macromolecule (= $4\pi R^2$ for spheres). The first term in Eq. 1.6 is the reversible work required to create a cavity of volume $\mathrm{V_s}$ within the polymer brush. The second term is the energy penalty associated with creation of a depletion layer around the colloid surface. The osmotic pressure term has been discussed earlier and is given by Eq. 1.5.

The interfacial tension for a planar interface is given by

$$\gamma(\phi) = -\Pi(\phi)\Gamma(\phi) + \int_0^{\phi} \Pi(\phi') \left(\frac{\partial \Gamma(\phi')}{\partial \phi'}\right) \, d\phi' , \tag{1.7}$$

where the relative adsorption $\Gamma(\phi)$ captures the effect of changes in local polymer density due to the creation of an interface. The adsorption is defined as

$$\Gamma(\phi) = \int_0^{\infty} \left(\frac{\phi(r)}{\phi(r \to \infty)} - 1\right) \, dr, \tag{1.8}$$

where $r$ is the distance from the colloid surface. The leading order expression of the adsorption is approximated by $\Gamma \approx -2R_g/\sqrt{\pi} \approx -R_g$. This assumes that the adsorption is independent of local density and behaves as an ideal polymer. With this approximation, the surface tension is described solely by the entropic free energy penalty of creating an additional cavity of volume $\Gamma \mathrm{A_s} \approx R_g \mathrm{A_s}$, in addition to the volume of the sensor $\mathrm{V_s}$.

Using this approximation, the surface tension term can be incorporated into a bulk osmotic pressure term in the insertion free energy as

$$\Delta U \approx \Pi(\phi) V^{\mathrm{eff}}, \tag{1.9}$$

where the effective volume of the cavity due to macromolecular insertion is $V^{\mathrm{eff}} = 4\pi R^3/3 + 4\pi R^2 R_g$ for a spherical polymer blob of size $R_g$ and macromolecule of size $R$ (see Fig. 1.2). For our system with $R \approx R_g$, the effective volume is well-approximated as $V^{\mathrm{eff}} = 4\pi(R + R_g)^3/3$.

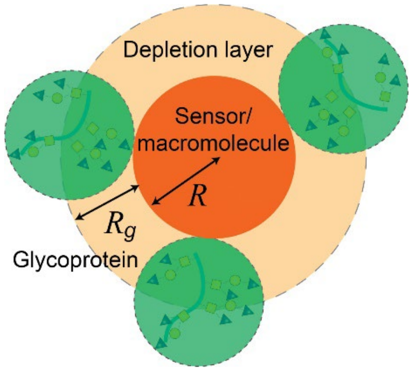


**Figure 1.2:** Schematic of the depletion layer and cavity related to the inclusion free energy of a colloid inside a polymer brush suspension. The size $R$ is the radius of the macromolecule and $R_g$ is the radius of gyration of the glycoprotein.

We use coarse-grained MD simulations to validate our theory. As shown in Fig. 1.3(A), when the insertion energy is plotted as a function of the surface density of polymers, there is no collapse of the data across different sensor sizes and lengths of the polymer brush. This indicates that the surface density is not an accurate, quantitative metric of crowding. Other proxy metrics like molecular weight and number density are inadequate to characterize crowding, because they do not include effects of variable glycoprotein length, polydispersity, stiffness, density, charge, and other interactions.

However, when the energy is plotted as a function of the osmotic pressure, the data collapse onto a master curve, as shown in Fig. 1.3(B). It is important to note that there is no adjustable fitting parameters here. These results confirm our hypothesis that the surface osmotic pressure is the unique quantity that provides a direct, quantitative metric of crowding. When quantifying crowding effects on cell surfaces, one should use the osmotic pressure instead of other proxy metrics like molecular weight, size, and surface density of the surface proteins.

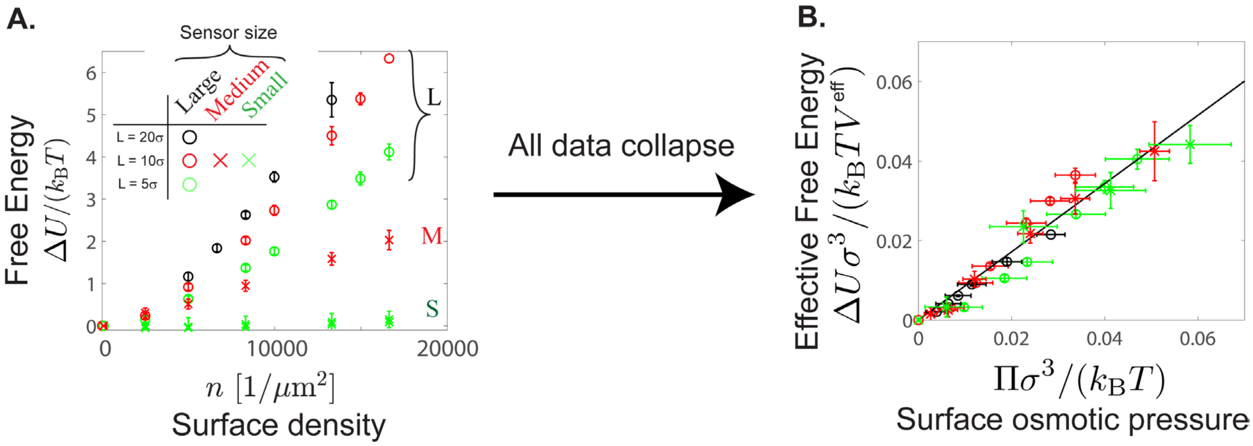


**Figure 1.3:** Insertion energy for large ($R = 2\sigma$), medium ($R = \sigma$), and small ($R = \sigma/2$) sensors as a function of surface polymer number density and contour lengths. (A) Insertion energy for the sensors as a function of surface density of polymers. The data do not collapse onto a universal curve, indicating that the surface density is not an accurate metric of surface crowding. (B) Normalized insertion energy, $\Delta U\sigma/(k_B T V^{\mathrm{eff}})$, where $V^{\mathrm{eff}} = 4\pi(R + R_g)^3/3$ is the excluded volume of the sensor in the polymer brush. Data across different sensor sizes and surface polymer contour lengths collapse onto a universal line when plotted as a function of osmotic pressure $\Pi$.

### 4. Analytical theory for effective brush potential

In our coarse-grained MD simulations, the Weeks-Chandler-Andersen (WCA) repulsive potential (10) was implemented between the polymer-polymer, polymer-sensor, polymer-substrate, and sensor-substrate. For the sensor-substrate interactions, we also add an attractive Morse potential to model affinity of the sensor to the membrane, which effectively approximates a short-range, attractive harmonic potential, $k(z-r_0)^2/2$, with stiffness $k$. These two potentials are sufficient to predict the free-energy landscape for the bare membrane simulations.

$$U_0 = \begin{cases} U_{\mathrm{WCA}} + U_{\mathrm{att}} & \text{if } z < 2^{1/6}R \quad (1.10) \\ U_{\mathrm{att}} & \text{if } 2^{1/6}R < z < (2^{1/6}R + R) \quad (1.11) \\ 0 & \text{if } z > (2^{1/6}R + R) \quad (1.12) \end{cases}$$

where the repulsive WCA potential is given by

$$U_{\mathrm{WCA}} = 4\epsilon\left[\left(\frac{R}{z}\right)^{12} - \left(\frac{R}{z}\right)^{6}\right] + \epsilon \tag{1.13}$$

and the attractive potential to the membrane surface is given by

$$U_{\mathrm{att}} = k(z-R)^2/2\ . \tag{1.14}$$

Obtaining the $\Delta U$ contribution to the overall free energy is non-trivial, since we only include WCA potentials for sensor-polymer interactions and no explicit potential of the brush into the sensor's equation of motion. We do not apriori know what the effective potential posed by the polymer brush will be.

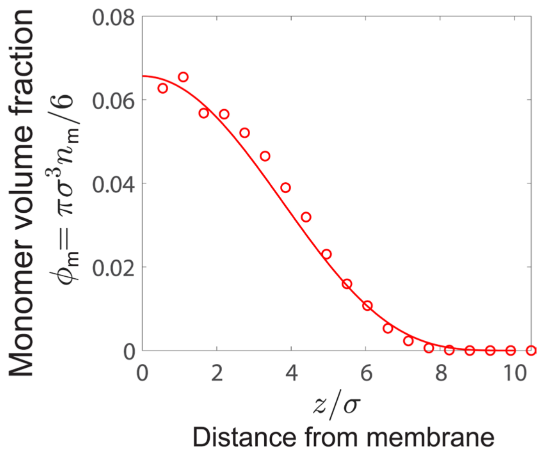


**Figure 1.6:** Local monomer volume fraction as a function of distance from the membrane, for a surface density of 8000 chains/$\mu$m$^2$. The symbols are simulation data, and the solid curve is Eq. 1.15.

From MD simulation data, the monomer volume fraction of the polymers as a function of height from the membrane surface is modeled by

$$\phi_m = \phi_m^0\left[1-\left(\frac{z}{L}\right)^2\right]^4. \tag{1.15}$$

We verified that this model for the polymer brush agrees with data from MD simulations, as shown in Fig. 1.6. The parabolic form inside the brackets is well known from polymer theory (11); we observe a steeper decay of density, likely due to the fact that the chains are not grafted and instead free to diffuse in 2D. The free energy of a sensor binding to a crowded polymer surface is given by

$$U = \begin{cases} U_{\mathrm{WCA}} + U_{\mathrm{att}} + \Delta U & \text{if } z < 2^{1/6}R \quad (1.16) \\ U_{\mathrm{att}} + \Delta U & \text{if } 2^{1/6}R < z < (2^{1/6}R + R) \quad (1.17) \\ \Delta U & \text{if } z > (2^{1/6}R + R) \quad (1.18) \\ 0 & \text{if } z > L \quad (1.19) \end{cases}$$

Substituting Eq. 1.15 into the virial pressure equation of state, Eq. 1.5, gives the local osmotic pressure as a function of distance from the membrane surface. The crowding energy as a function of distance, $\Delta U(z)$, is obtained by using Eq. 1.9. The resulting crowding energy is used in Eqs. 1.16 – 1.19 to obtain the total energy as a function of distance, $U(z)$. As shown in Fig. 1.7, the data from our MD simulations (black symbols) demonstrate excellent agreement with our theory (red curve).

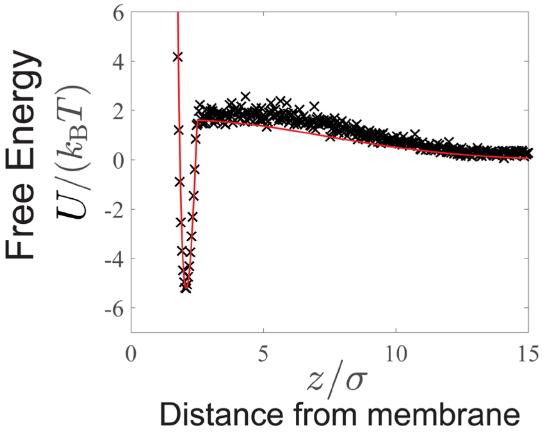


**Figure 1.7:** Full free energy curve using Eqs. 1.16 - 1.19 (solid red curve) agrees with the direct calculation from the sensor probability distribution (black crosses). There are no adjustable parameters in the theoretical model, Eqs. 1.16 – 1.19.

### Chapter II. Coarse-grained molecular dynamics (MD) simulations

#### 1. Basic construction of the model

To construct a molecular model of macromolecular transport across cell surface proteins and glycocalyx, we performed coarse-grained Molecular Dynamics (MD) simulations of colloidal transport within semi-flexible polymers diffusing on 2D surfaces. This is a modified extension of our protein polymer surface model presented previously (12).

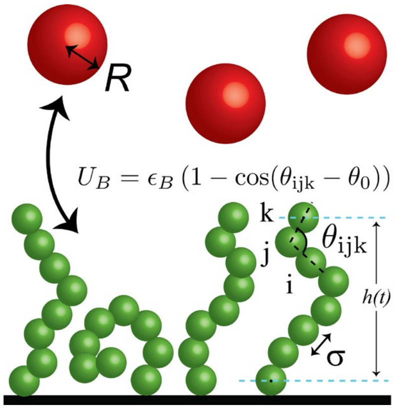


**Figure 2.1:** Schematic of our coarse-grained MD simulations of a protein polymer brush with surface-binding macromolecular sensors. Using a Kremer-Grest bead-spring model (13), individual beads of diameter $\sigma$ are connected by a finitely extensible nonlinear elastic (FENE) potential and a bending stiffness is invoked by a 3-particle angular potential, $U_B$. The bottom bead is confined to remain along a 2D surface but can diffuse laterally along the surface. Sensor particles of size $R$ are added to the bulk above the polymer brush. The sensors have a short-ranged binding affinity to the substrate.

We modeled surface protein chains using a Kremer-Grest bead-spring model (13), with each bead representing a structured protein domain or a coarse-grained unit of an intrinsically disordered domain. A bead at one end of the chain is confined between two parallel walls separated by one bead diameter, allowing it to diffuse freely in 2D but cannot escape out of plane. All other beads on the chain are free to move in 3D, except through a bottom wall that acts as a solid substrate, thereby modeling protein diffusion along a membrane that is in-plane fluid. Because the size of each protein domain is large compared to the surrounding solvent molecules, the solvent is coarse grained and its dynamics are not explicitly evolved. In other words, the protein chain experiences a hydrodynamic drag and Brownian motion from the continuous solvent. In this work, the membrane does not deform nor fluctuate out of plane, although these effects may be included.

Simulations were performed using a GPU-enabled HOOMD-blue molecular dynamics package (14,15), and all simulations contained at least 2000 protein chains and 10000 sensor particles. The dynamics of particle $i$ is evolved in time following the overdamped Langevin equation

$$\mathbf{0} = -\zeta_i \boldsymbol{U}_i + \boldsymbol{F}_i^B + \boldsymbol{F}_i^P, \tag{2.1}$$

where $\zeta_i$ is the hydrodynamic drag factor, $\boldsymbol{U}_i$ is the velocity, $\boldsymbol{F}_i^B = \sqrt{2\zeta_i^2 D_i}\boldsymbol{\Lambda}$ is the translational Brownian force, $D_i$ is the Stokes-Einstein-Sutherland translational diffusivity of a single monomer, and $\boldsymbol{F}_i^P$ is the interparticle force. The left-hand side is zero since inertia is negligible for coarse-grained proteins embedded in a viscous solvent. The translational diffusivity is modeled with the usual white noise statistics, $\langle \boldsymbol{\Lambda}(t) \rangle = \mathbf{0}$ and $\langle \boldsymbol{\Lambda}(t)\boldsymbol{\Lambda}(0) \rangle = \delta(t)\, \boldsymbol{I}$, where $\delta(t)$ is a delta function and $\boldsymbol{I}$ is the identity tensor. The drag coefficient $\zeta_i$ of particle $i$ is linearly scaled with the particle size $\sigma_i$. The interparticle force (described below) includes contributions from sensor-polymer chain interactions, intrachain polymer potential, interchain bead-bead pair interactions, and bead-wall interactions.

All interactions between the particles and walls are modeled with a Weeks-Chandler-Andersen (WCA) potential (10), in which a Lennard-Jones (LJ) potential is shifted upwards, truncated at the potential minimum of $2^{1/6}\,\sigma_i$ (such that the potential is purely repulsive), and assigned a well depth of $\epsilon = k_B T$ (where $k_B T$ is the thermal energy). All lengths are expressed in units of the LJ diameter $\sigma$ and is set to unity. We connect the polymer chains with a finitely extensible nonlinear elastic (FENE) potential, using a spring constant of $k_0 = 30$ and a bond length of $R_0 = 1.5$ (expressed in terms of reduced LJ units, $\epsilon = \sigma = 1$). To model semi-flexible polymers, we implemented a bending potential between 3 neighboring particles to capture chain stiffness, $U_B = \epsilon_B\big(1 - \cos(\theta_{\mathrm{ijk}} - \theta_0)\big)$, where $U_B$ is the bending energy, $\theta_{\mathrm{ijk}}$ is the bond angle between neighboring particles ($i,j,k$), and $\theta_0 = \pi$ is the resting angle. The persistence length of the chains was measured by calculating the bond angle correlation, $\langle \boldsymbol{e}_1 \cdot \boldsymbol{e}_i \rangle = \exp(-s_i/\ell_P)$, where $\boldsymbol{e}_i$ is the unit vector connecting the center of mass positions of particles $i$ and $i+1$, and $s_i$ is the path length along the polymer to particle $i$. To implement the persistence length $\ell_P$ as an input to the simulations, the bending energy is set to $\epsilon_B/(k_B T) = \ell_P/\sigma$ to achieve the desired polymer stiffness. We have set $\ell_P = 3\sigma$ for all of our protein-based simulations based on previous work, and have verified the angular correlation for unbound polymers in 3D as a validation of proper implementation (12).

Sensor particles of different sizes were added to the bulk of the simulation box to model the dynamics of sensor transport and binding to the cell surface. The diameters of the Chol-0.5k PEG, 10k dextran, and 40k dextran sensors were modeled with spheres of diameter 3, 5, and 10 nm, respectively (16). The sensor particles have an attractive potential to the membrane via a Morse potential with the HOOMD parameters ($D_0 = 3 \times 10^9$, $\alpha = 10^{-4}$, $r_0 = \sigma/2$, and $r_{\mathrm{cut}} = \sigma/2 + R$), which effectively approximates a short-range, attractive harmonic potential $U_{\mathrm{att}} = k(z - r_0)^2/2$ with stiffness $k = 2D_0\alpha^2 = 60$. All other particle pairs experience a short-ranged repulsive WCA potential, as described earlier. The dynamics of sensor particle $i$ is evolved in time following the Langevin equation

$$\mathbf{0} = -\zeta_i \boldsymbol{U}_i + \boldsymbol{F}_i^B + \boldsymbol{F}_i^P + \boldsymbol{F}_i^{\mathrm{wall}}, \tag{2.2}$$

where $\boldsymbol{F}_i^{\mathrm{wall}} = \boldsymbol{F}_i^{\mathrm{att}} + \boldsymbol{F}_i^{\mathrm{WCA}}$ is the short-ranged potential that the sensor particle experiences with the surface, $\boldsymbol{F}_i^{\mathrm{att}} = -\nabla U_{\mathrm{att}}$ is the attractive linear force, and $\boldsymbol{F}_i^{\mathrm{WCA}} = -\nabla U_{\mathrm{WCA}}$ is the WCA potential with the surface (see

Eqs. 1.13 and 1.14 for definitions of the potentials). In summary, each sensor particle experiences the potentials in Eqs. 1.10 – 1.12 as a function of height $z$, in addition to the pair-wise WCA potentials with the protein polymers. It is straightforward to add flexibility and non-spherical shape effects into our sensor model.

Unlike our previous work (12) where a large concentration of passive PEG depletants were added to the bulk to compress the cell surface proteins, in this work we focus on dilute concentrations in which the osmotic compression is negligible. Furthermore, we did not observe any depletion flocculation nor any large-scale protein clustering. Simulations with dilute concentrations of sensors are very time consuming because acquiring sufficient statistics requires many time steps at equilibrium. Therefore, we accelerated our simulations by using a larger concentration of sensor particles that interact only with the protein polymer and the surface, but not with each other. The sensors act as "ideal-gas" particles to each other but interact with WCA potentials with the surface proteins. However, sensor concentrations that are too large can interact with each other indirectly via correlated motion of the protein polymers. A sensor volume fraction less than 1% was used in this work.

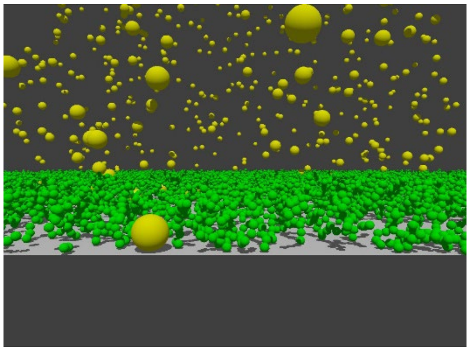

**Figure 2.2:** Snapshot of coarse-grained MD simulations with sensors (yellow particles) of size $R$ = 2$\sigma$, and surface polymers (green particles) of length $L$ = 10$\sigma$, where $\sigma$ is the size of an individual monomer.

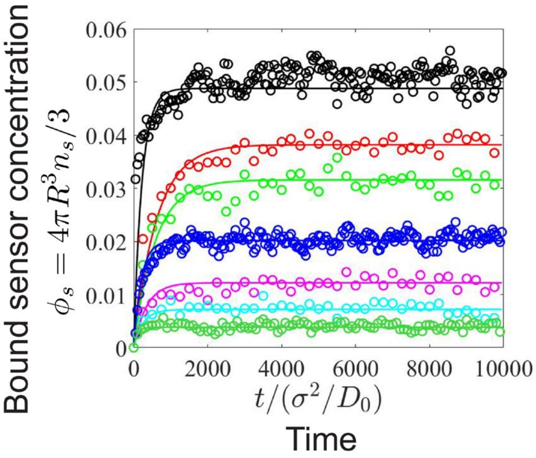


**Figure 2.3:** Volume fraction of sensors of size $R = \sigma$ on the membrane surface, $\phi_s = 4\pi R^3 n_s/3$, as a function of nondimensional time $t/(\sigma^2/D_0)$, where $D_0$ is the Stokes-Einstein-Sutherland diffusivity based on the monomer size $\sigma$. Solid curves are a fit to a first-order kinetic rate law, $\phi_s/\phi_s^\infty$=1 - exp(-t/$\tau$).

We used a system box size of $V = L^2 L_z$, where $L$ was adjusted to achieve the specified area density and number of polymer chains, and $L_z$ was chosen to be sufficiently large to maintain a uniform bath concentration of sensors in the bulk. Dilute surface densities were conducted with ~ 200 chains/$\mu$m$^2$, and the dense surfaces contained ~ 20000 chains/$\mu$m$^2$. We imposed periodic boundary conditions in the $x$ and $y$ directions, and no-flux hard walls at $z$ = 0 and $z = L_z$ to prevent any particles from passing through. Initial configurations were generated by placing the particles in lattice locations, and sufficient time steps were run to reach a steady-state. Time steps were varied from $\Delta t = 10^{-6} - 10^{-4}$ and verified to be sufficiently small to capture relevant dynamics.

A snapshot of the simulations is shown in Fig. 2.2. Simulations were conducted for long enough time steps to sample equilibrium properties. Figure 2.3 shows the concentration of sensors on the membrane surface as a function of time, and we can see that sufficient configurations are sampled after equilibrium is achieved.

### 2. Calculation of mechanical stress (and pressure) of cell surface proteins

The pressure of cell surface proteins is given by the negative trace of the virial stress tensor,

$$\boldsymbol{\sigma} = -nk_BT\boldsymbol{I} - \frac{1}{V}\langle \sum_i \sum_{j>i} \boldsymbol{r}_{ij}\boldsymbol{F}_{ij}^P \rangle, \tag{2.3}$$

where $V = L^2\langle h \rangle$ is the volume of the polymer brush, $L$ is the length of the simulation box, $\langle h \rangle$ is the average height of the polymer surface, $\boldsymbol{r}_{ij}$ is the interparticle distance, and $\boldsymbol{F}_{ij}^P$ is the pair-wise interparticle force between all particles, including individual monomers within a single polymer. The first term in Eq. 2.3 is the ideal-gas stress and the number density is defined as $n = N/V$ where $N$ is the total number of particles. The second term is the Irving-Kirkwood virial stress tensor that measures the contribution from interparticle interactions. For polymers, it is important to note that the ideal-gas pressure is $\Pi^{\mathrm{ig}} = n_c k_{\mathrm{B}}T$ and not $nk_{\mathrm{B}}T$, where $n_c$ is the number density of chains, $n_c = N_c/V$ ($N_c$ is the total number of chains). Particles lose their translational degree of freedom when constrained as a polymer, so the ideal-gas pressure decreases with increasing degree of polymerization. Returning to our virial EOS in Eq. 1.5, we can rewrite the ideal-gas expression as $\Pi^{\mathrm{ig}} = (k_{\mathrm{B}}T/V_p)\phi/N_R$, where the degree of polymerization $N_R = N/N_c$. For very large molecular weight polymers ($N_R \rightarrow \infty$), the ideal-gas contribution is zero and the first term contributing to the osmotic pressure is two-body interactions (i.e., second virial coefficient).

In calculating the osmotic pressure using Eq. 2.3, it would seem like we are overestimating the pressure based on the first term, which assumes that all $N$ particles are independent. However, the virial stress term will cancel the "excess" stress contribution from the $nk_{\mathrm{B}}T$ term. That is, the FENE potential applied to the constitutive particles on the polymers (which represent elasticity) has the opposite sign and subtracts the independent degrees of freedom. The net result is that we correctly recover the ideal-gas pressure of a polymer, $\Pi^{\mathrm{ig}} = (k_{\mathrm{B}}T/V_p)\varphi/N_R$.

### 3. Calculation of free energy potentials

The free energy experienced by the sensor particle as a function of the distance from the membrane surface is given by $U = -k_{\mathrm{B}}T \ln P(z)$, where $P(z)$ is the normalized probability distribution of the sensors. We calculate $P(z)$ by binning the simulation into thin slabs and ensemble averaging over all particles and time. From these distributions, we calculate the effective binding energy and the insertion penalty at equilibrium, as shown in Fig. 2.4. The offset at the energy minimum gives the glycocalyx contribution $U_g$ and the in-plane osmotic pressure of the cell surface.

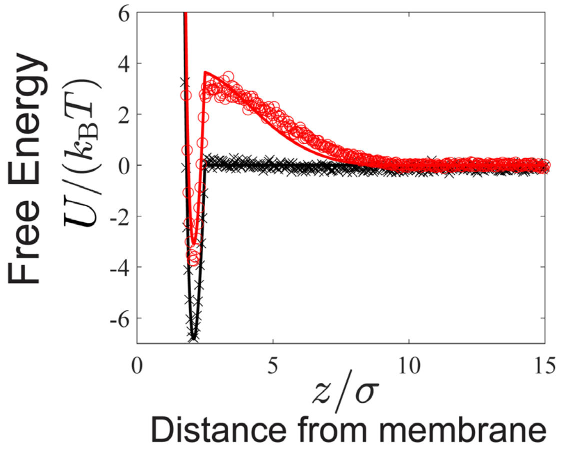


**Figure 2.4:** Effective free energy potential experienced by a macromolecule of size $R = \sigma$ approaching a bare (black) and crowded (red) cell surface membrane, which is a superposition of two contributions, $U_0$ and $\Delta U$. The circle and cross symbols are a direct calculation of the free energy via $U = -k_{\mathrm{B}}T \ln P(z)$ from

MD simulations, where $P(z)$ is the normalized probability distribution of the sensors. Solid curves are Eqs. 1.16-1.19. The depth of the minimum gives the effective binding energy at equilibrium.

As shown in Fig. 2.4, our analytical theory in Eqs. 1.16-1.19 agrees very well with the direct energy calculations from MD simulations. Once the sensor-substrate affinity is determined in the theory by fitting to the bare-membrane simulations (black symbols and curve), there are no fitting parameters for the crowded surface simulations and theory (red symbols and curve).

#### 4. Incorporation of charges on protein polymers

Charge effects were analyzed in MD simulations by adding a screened coulomb (Yukawa) pair potential between the constituent monomers of the polymer. The energy scale was set to $\epsilon = k_{\mathrm{B}}T$, and the Debye length was varied from $\kappa^{-1} = \sigma$ to $\kappa^{-1} = 2\sigma$. Electrostatic interactions were not included for sensor-polymer or sensor-sensor interactions.

### Chapter III. Experimental details

#### 1. Control experiments of polymer-based sensors with different chemistries.

Our dextran-based sensors have a strong affinity to insert into the lipid bilayer via cholesterol tags bound to the dextran molecules. To rule out the possibility of other, non-specific interactions of the dextran-based sensors onto membranes, we conjugated the dextran molecules to Alexa Fluor dyes only, without cholesterol tags. Upon incubating these dextran-dye sensors with lipid-coated beads and red blood cells (RBCs), we did not detect any binding onto the membrane, validating that the main adsorption mechanism of our sensors is via the cholesterol tags.

We further validated that the negative charges present on Alexa Fluor dyes do not impact sensor binding onto cell surfaces. In order to test whether the negative charges on the Alexa Fluor dyes interact with the sialic acid on the red blood cell surface, we synthesized dextran sensors using BODIPY, a charge-neutral dye. We observed no difference in binding between the Alexa Fluor and BODIPY conjugated sensors. We conclude that our sensors interact only sterically with the crowders on the cell surface. Negative charges on the cell surface generate intrachain and interchain stretching of the surface proteins, but the surface polymers do not interact electrostatically with our sensors.

#### 2. Verification that the stock sialidase is protease-free.

A sodium dodecyl sulfate polyacrylamide gel electrophoresis (SDS-PAGE) protein gel of bovine serum albumin (BSA), MW = 66 kDa, is shown in Fig. 3.1. BSA was treated with various concentrations of sialidase and Proteinase K at 37C for 5 hours. The protein was subsequently heat-denatured in 1x Laemmli Sample Buffer (Sigma Aldrich) in the presence of $\beta$-mercaptoethanol. The sample was then loaded onto a NuPAGE Novex 4–12% gradient Bis-Tris gel (Fisher Scientific) and separated by electrophoresis. This gel confirms that our stock sialidase does not have protease activity. In addition to BSA, we collected the supernatant of RBCs treated with sialidase and found no detectable signal of proteins released from the RBC surface as a result of sialidase treatment. This verifies that soluble sialic acid-binding proteins are not embedded in the glycocalyx to hinder the binding of the sensors.

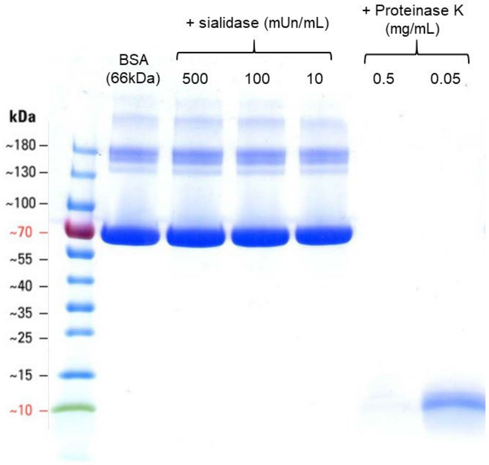


**Figure 3.1**. Sodium dodecyl sulfate polyacrylamide gel electrophoresis (SDS-PAGE) protein gel of bovine serum albumin (BSA), MW = 66 kDa, treated with enzymes sialidase and Proteinase K at 37C for 5 hours. We detected no protease activity in our stock sialidase. In our experiments, RBCs were treated with sialidase at 50 mUn/mL.

### 3. EDC conjugation to remove negative charges from cell surface

We used a carbodiimide crosslinker chemistry based on 1-Ethyl-3-(3-dimethylaminopropyl)carbodiimide (EDC) to remove negative charges present on carboxylic acid groups on the red blood cell surface. Most surface negative charges come from the carboxylic acid on the sialic acids, but charged amino acids (e.g., aspartic acid and glutamic acid) may also be impacted by this reaction. We added EDC and hydrazide-biotin (a charge-neutral molecule) to a suspension of red blood cells, and the effectiveness of the reaction was assessed by imaging with AF555-labeled straptavidin. As shown in Fig. 3.2, the reaction was qualitatively very effective and we believe that most negative charges were neutralized on the cell surface.

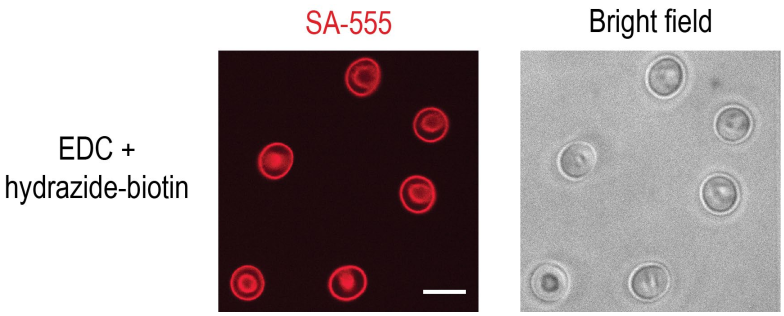


**Figure 3.2**. EDC chemistry was used to remove negative charges from the red blood cell surface. The reaction conjugated hydrazide-biotin to carboxylic acid sites, which enabled detection using AF555-labeled streptavidin to image the red blood cell surface. Scalebar is 10 $\mu$m.
